# Placenta DNA methylation at *ZNF300* is associated with fetal sex and placental morphology

**DOI:** 10.1101/2021.03.05.433992

**Authors:** Christine Ladd-Acosta, Shan V. Andrews, Kelly M. Bakulski, Jason I. Feinberg, Rakel Tryggvadottir, Ruofan Yao, Lisa A. Croen, Irva Hertz-Picciotto, Craig J. Newschaffer, Carolyn M. Salafia, Andrew P. Feinberg, Kasper D. Hansen, M. Daniele Fallin

**Affiliations:** Department of Epidemiology, Johns Hopkins Bloomberg School of Public Health, 615 N. Wolfe Street, Baltimore, MD 21205, USA; Wendy Klag Center for Autism and Developmental Disabilities, Johns Hopkins Bloomberg School of Public Health, 615 N. Wolfe Street, Baltimore, MD 21205, USA; Department of Epidemiology, University of Michigan School of Public Health, 1415 Washington Heights, Ann Arbor, MI 48109, USA; Department of Mental Health, Johns Hopkins Bloomberg School of Public Health, 624 N. Broadway, Baltimore, MD, 21205, USA; Center for Epigenetics, Institute for Basic Biomedical Sciences, Johns Hopkins School of Medicine, 733 N. Broadway, Baltimore, MD 21205, USA; Department of Obstetrics and Gynecology, Loma Linda University School of Medicine, 11234 Anderson St, Loma Linda, CA 92354, USA; Division of Research, Kaiser Permanente Northern California, 2000 Broadway, Oakland, CA 94612, USA; Department of Public Health Sciences, School of Medicine, University of California Davis, 4610 X St, Sacramento, CA 95817, USA; MIND Institute, University of California Davis, 2825 50^th^ St, Sacramento, CA 95817, USA; AJ Drexel Autism Institute, Drexel University, 3020 Market St #560, Philadelphia, PA 19104, USA. Present address: The Pennsylvania State University, College of Health and Human Development, 325 Health and Human Development Building, University Park, PA, 16802, USA; Department of Epidemiology and Biostatistics, Drexel University Dornsife School of Public Health, 3125 Market St, Philadelphia, PA 19104, USA; Placental Analytics LLC, 187 Overlook Circle, New Rochelle, NY 10984, USA; Department of Medicine, Johns Hopkins School of Medicine, 733 N. Broadway, Baltimore, MD 21205, USA; Department of Genomic Medicine, Johns Hopkins School of Medicine, 733 N. Broadway, Baltimore, MD 21205, USA; Department of Biostatistics, Johns Hopkins Bloomberg School of Public Health, 615 N. Wolfe Street, Baltimore, MD 21205, USA

## Abstract

Fetal sex-specific differences in placental morphology and physiology have been associated with sexually dimorphic health outcomes. However, the molecular mechanisms underlying these sex differences are not well understood. We performed whole genome bisulfite sequencing in 133 placenta samples and discovered a significant difference in DNA methylation (DNAm) at the *ZNF300* gene locus between male and female offspring and replicated this result in 6 independent datasets. Additionally, the sex-specific pattern appears to be placenta-specific, is robust to a wide range of gestational ages and adverse health outcomes and is present in sorted placenta villous cytotrophoblast cells. Integration of DNAm, genetic, and placental morphology data from the same individuals revealed *ZNF300* methylation is also associated with placenta area, perimeter, and max diameter, genetic variants on chromosomes 5 and X, and may mediate the effects of genetic variation on placental area.

## Introduction

The placenta is a vital organ that is only present in women during pregnancy. It plays a critical role in sustaining pregnancy and supporting fetal growth and development. Additionally, it regulates the *in*-*utero* environment, which has been shown to influence health outcomes across the lifespan[1-3]. Placental size, morphology, vascular structures, and physiologic processes have been associated with a wide range of adverse health outcomes in pregnant women[4, 5] and their offspring’s outcomes at birth[6-11], in childhood[12-15], in adolescence[16], and in adulthood[16-22]. Furthermore, sex-specific differences in placenta morphology and physiology have been reported[23-28] and linked to sexually dimorphic health outcomes and/or response to environmental cues[29-31]. However, the molecular mechanisms underlying sex-specific differences in placental growth, function, and response to the *in-utero* environment are not well understood.

Genetic variation may help this understanding. Sex-specific differences in the effects of genetic risk factors on health outcomes have been reported[32] including for mortality rates after injury [33], susceptiblity to immune diseases [34], response to vaccination [35], chronic obstructive pulmonary disease risk[36], basal cell carcinoma[37], cardiometabolic disorders [38, 39], and depression [40, 41]. Genetic variation in placental tissue has also been shown to impact birth outcomes including large for gestational age [42], birthweight [43, 44], and preterm birth [45], but studies to investigate differences in genetic risks by sex or to identify genetic variant associations with placenta size and morphology are lacking. To our knowledge, only 2 studies have examined single nucleotide polymorphism associations with inter-individual differences in placental growth characteristics. The first was a candidate gene study in which the authors reported associations between genetic variation at the *CYP2A6* gene locus, placental weight, and newborn birth weight [46]. The second reported differences in placental weight with increasing aggregate polygenic risk for newborn birth weight[45]. Given the paucity of research in this area, additional studies are needed to identify genetic variants associated with placental size and to assess potential differences in their effects by fetal sex.

Epigenetic mechanisms could also provide molecular mechanisms to explain sex-differences in placental morphology, function, and fetal birth outcomes. Gene expression differences in placenta related to fetal sex have been observed [47-49] and some have been shown to mediate genetic risk effects on birthweight [44]. In addition to gene expression, changes in DNA methylation have also been observed. Locus-specific differences in DNA methylation by sex have been observed in human peripheral blood[50-52], cord blood[53], pancreas[54], and prefrontal cortex[55] tissues. In addition, studies have identified locus-specific methylation changes in placenta related to fetal sex [56], and sexually dimorphic associations between placenta DNA methylation and environmental toxicants [57-59], sociodemographic factors[60], maternal conditions[61-65] and offspring health outcomes[66, 67]. Sex-specific differences in the epigenetic aging pathway[68] and aggregate genomic DNA methylation levels [69] have also been observed in placental samples. While previous findings support the potential for DNA methylation to play a role in sex-specific differences in placenta and associated environmental exposures and health outcomes, they have been limited by small sample sizes [54, 56] or examination of a small proportion of the genome (∼2% of methylome)[50-54, 68] or have been limited to non-developmental tissues[53, 54]. Furthermore, no studies have evaluated whether differences in placenta DNA methylation patterns, by sex, are also associated with differences in placenta size or genetic variation.

Here, we expand upon previous findings by performing a whole genome scan in a moderately sized sample and identified a novel significant difference in DNA methylation pattern by fetal sex at the zinc finger protein 300 (*ZNF300*) gene locus. We performed replication analyses in samples from six independent studies, at different gestational windows, in a specific isolated cell type, and across a variety of health outcomes, demonstrating the robustness of our result. Finally, we assessed whether the sex-specific DNA methylation patterns we identified at *ZNF300* were associated with measures of placental size and genetic variants to better understand whether DNA methylation mediates the effects of genetic variation on placental size and in a sex-specific manner.

## Results

### Whole-genome scan to identify placenta methylation differences associated with fetal sex

Whole genome bisulfite sequencing data were obtained for a total of 133 placentae collected from participants enrolled in the Early Autism Risk Longitudinal Investigation (EARLI) cohort. As shown in Supplementary Table 1, no significant differences in gestational age, self-reported race, mode of delivery, or loss to follow up were found between male and female offspring. However, a larger number of male offspring went on to receive an autism spectrum disorder diagnosis at age 3, as expected. For our initial discovery screen - to identify differential methylation in placenta related to fetal sex - we only included placentae from males and females with typical neurodevelopment to address this difference as well as any potential concerns that differences could be related to autism diagnoses. Thus, our genome-wide discovery screen included 37 total placentae including 17 from male offspring and 20 from female offspring with neurotypical development. We identified a single DMR that was significantly associated (p_permutation_ = 0.015) with fetal sex after applying a permutation-based method to correct for multiple testing and controlling for a family-wise error rate (FWER) using a < 0.05 threshold. The DMR showed a 15% increase in DNA methylation, on average, in placentae from males compared to females and is located in a 1597 bp region on chromosome 5 in the CpG island promoter region of zinc finger protein 300 (*ZNF300*). It contains placenta specific DNaseI hypersensitive sites (Fig. 1). We observed no differences in sequencing coverage of this region related to fetal sex (Fig. 1b). We also investigated whether the DNA sequence of the *ZNF300* DMR shared any degree of homology to DNA sequence present on the sex chromosomes; no sex chromosome sequence homology was found (Supplementary Table 2). Finally, no significant differences in gestational age, race, or mode of delivery were related to sex in the overall set of EARLI placenta samples (Supplementary Table 1), nor were these factors related to DNA methylation at *ZNF300* (Supplementary Table 3). A full list of candidate DMRs we identified at suggestive p value thresholds is provided in Supplementary Table 4, and a volcano plot showing the results of our genome-wide screen is provided in Supplementary Fig. 1.

**Figure 1:**
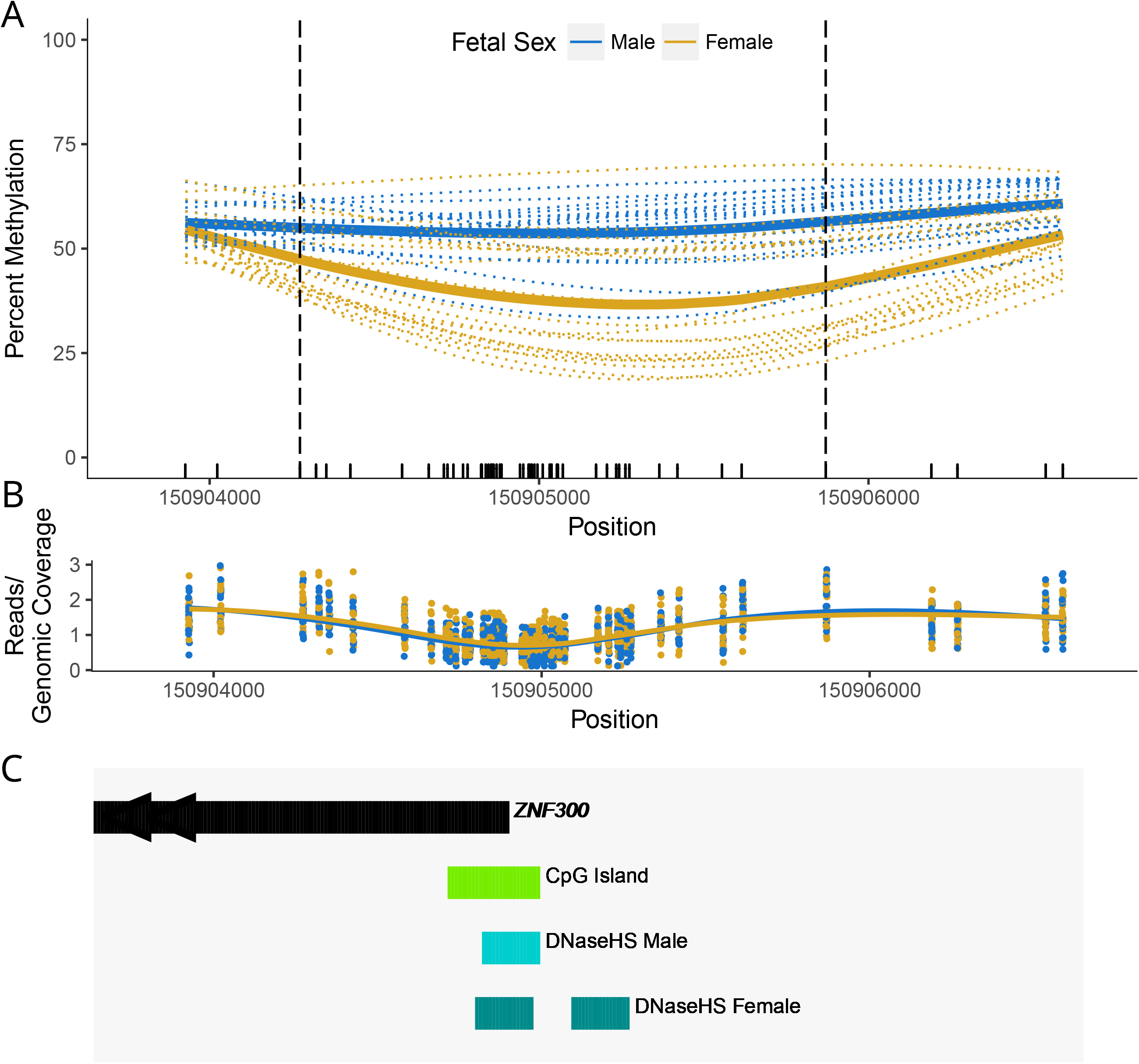
Placenta methylation levels, at *ZNF300*, are associated with by fetal sex. Samples from male and female offspring are denoted in blue and yellow, respectively. **A**, Percent methylation (y-axis) plotted against genomic position (x-axis). Dashed horizontal lines represent individual samples and solid lines represent average s across all male or female samples. Vertical dashed lines denote differentially methylated region (DMR) start and end position. Each tick mark at the bottom of the plot represents a CpG site. **B**, Sequencing read counts (divided by average coverage across all autosomal loci; y-axis) vs. genomic position (x-axis). **C**, Gene and genomic feature annotation, obtained from the UCSC Table Brower and ENCODE databases.

### Replication and robustness of fetal sex-specific placenta DNA methylation levels at *ZNF300*

First, we evaluated whether an independent set of placentae from EARLI participants that went on to have atypical neurodevelopment, including 26 from male and 24 from female offspring, also showed sex-related DNA methylation differences at *ZNF300*. Consistent with our discovery results, we observed higher DNA methylation levels, 12% on average, in placenta from women carrying male compared to female offspring (Fig. 2a). We also examined DNA methylation measures from six additional datasets that use a highly reliable, independent, measurement platform – the Illumina 450K BeadChip (450K). A summary of the replication datasets we used for these purposes is provided in Supplementary Table 5. We identified 10 probes measured on the 450K that overlapped the physical position of the sex-associated differentially methylated region (DMR), at *ZNF300*, that we identified using whole-genome bisulfite sequencing. In all 6 datasets, we observed higher DNA methylation levels in placentae from male offspring compared to females, ranging in magnitude from 12% to 22% (Fig. 2b-g), across this genomic region. The increase in DNA methylation among males was present in both term and preterm placenta, including those from first and second trimester pregnancies (Fig. 2f-g). Furthermore, we found evidence that the DNA methylation differences by offspring sex are not simply reflecting differences in cell type proportions from heterogenous placentae samples. Examination of a single placenta cell type, villous cytotrophoblasts, also showed increased DNA methylation among males (Fig. 2g), with the largest magnitude of sex difference in DNA methylation (25%) across the replication data sets. Several of the replication datasets included samples from women that had pregnancy complications (PR4, PR5, PR6) or whose children went on to have atypical neurodevelopment in childhood (PR1). The sex-related differences at *ZNF300* were robust across these adverse health outcomes (Fig, 2a, d-f).

**Figure 2:**
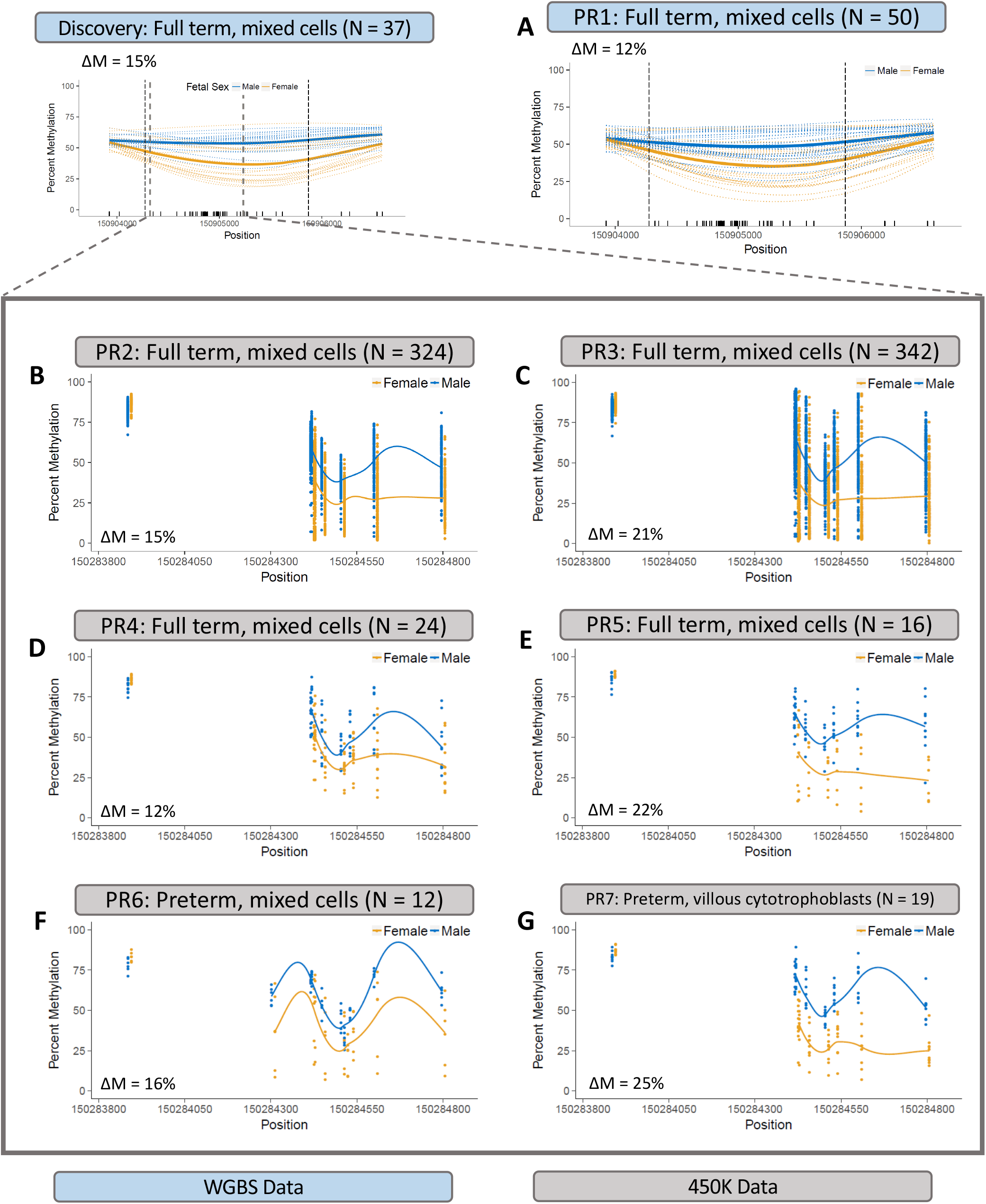
Replication of sex-specific differential placenta methylation at *ZNF300*. For each panel, percent methylation (y-axis) is plotted against genomic position (x-axis). Dashed horizontal lines represent individual samples and solid lines represent average s across all male or female samples. Vertical dashed lines denote differentially methylated region (DMR) start and end position. Tick marks at the bottom of the plot represent CpG sites. Samples from male and female offspring are denoted in blue and yellow, respectively. ΔM stands for difference in methylation between males and females. **A**, Whole genome bisulfite sequencing (WGBS) methylation measures in a set of Early Autism Risk Longitudinal Investigation samples with atypical neurodevelopment that were not included in the discovery phase. Replication datasets with Illumina 450K placenta methylation measures from: **B-C**, term pregnancies with bulk placenta tissue; **D-E**, term pregnancies with bulk placenta tis sue and maternal pregnancy complications; **F**, bulk placenta tissue from preterm pregnancies with maternal complications; **G**, preterm pregnancies with sorted villous cytotrophoblast cells. Note: 450K array probes only cover a portion of the DMR discovered with WGBS data.

Finally, we evaluated whether the sex-specific differential methylation pattern at *ZNF300* was also present in other developmentally relevant biospecimens including cord blood (n = 223), fetal brain (n=179), and childhood peripheral blood (n=970). As shown in Fig. 3, we did not observe the same large magnitude of change in DNA methylation between males and females for any of the other 3 developmental samples.

**Figure 3:**
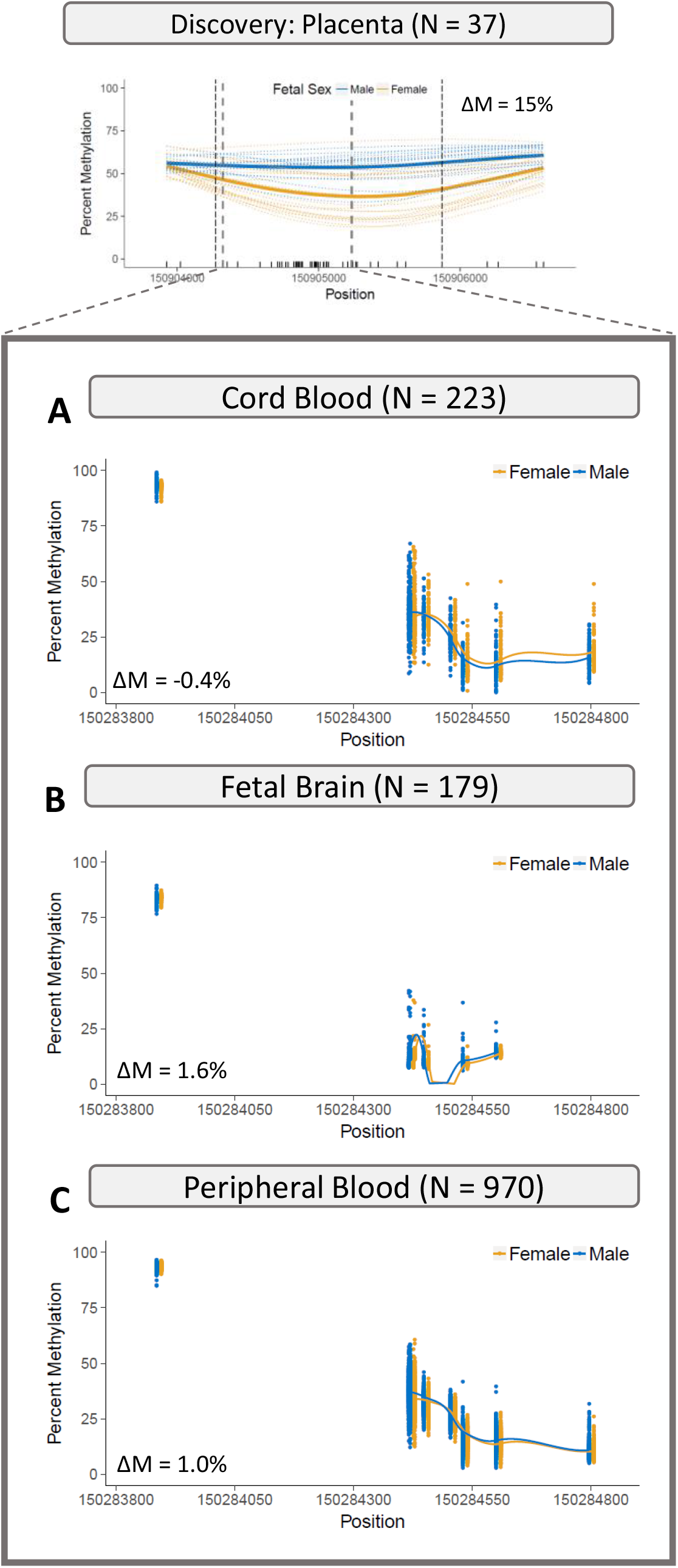
Tissue-specificity of sex-specific methylation changes at *ZNF300*. Percent methylation (y-axis) is plotted against genomic position (x-axis). Points represent individual samples and smoothed lines represent methylation average, across all male (blue) or female (yellow) samples. DNA methylation data obtained using the 450K array in **A**, cord blood samples from the EARLI study; **B**, fetal brain samples; **C**, peripheral blood in early childhood.

### *ZNF300* methylation associations with fetal growth

Increases in DNA methylation in the promoter region of the *ZNF300* locus were previously associated with intrauterine growth restriction (IUGR) among 8 monochorionic IUGR discordant twin pair samples[67]. We sought to replicate the finding that methylation at the ZNF300 locus is related to fetal growth in an independent sample and also evaluate whether there were sex-specific differences in the relationship between DNA methylation and fetal growth. In the EARLI whole-genome bisulfite data (n=74), we observed no differences in *ZNF300* methylation related to newborns being small for their gestational age (SGA; Supplementary Fig. 2a). However, among male offspring, newborns that were SGA had less methylation than those that were not SGA (Supplementary Fig. 2b). Interestingly, the opposite pattern was observed in female newborns. Females that were SGA showed an increase in DNA methylation, on average, compared to those without SGA (Supplementary Fig. 2c) at this locus. In a second publicly available dataset (GSE71678) with 450K and SGA data available, we observed similar decreased methylation related to SGA compared to non-SGA in male offspring placenta but no change in methylation related to SGA among females (Supplementary Fig. 2d-f).

### *ZNF300* methylation levels are associated with placenta morphological features

We evaluated the relationship between methylation levels at the *ZNF300* locus and placenta morphological features, specifically the placental perimeter, area, and max diameter (determined by 2D fetal surface imaging) using EARLI samples with both methylation and morphology data available (N = 53). As shown in Fig. 4, the average methylation level across the *ZNF300* DMR was significantly associated with placenta perimeter (p = 0.036), area (p = 0.014), and max diameter (p = 0.037), adjusted for feto-placental weight ratio and gestational age. Sex-stratified analyses revealed that this association was driven by female samples; no association were observed in males (Supplementary Fig. 4).

**Figure 4:**
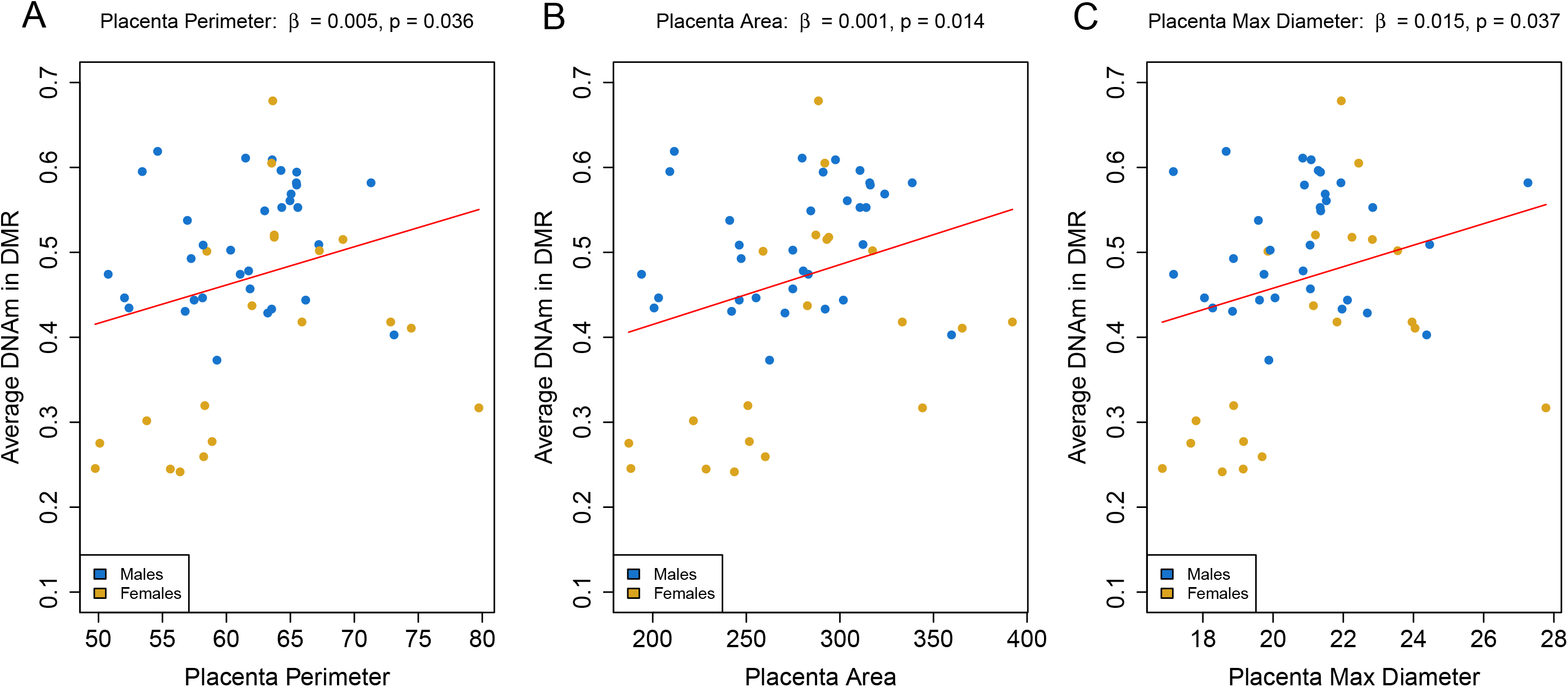
Relationship between DNA methylation level at the *ZNF300* locus and placenta size measures. Mean DNA methylation levels (y-axis) plotted against placenta: **A**, perimeter (cm); **B**, area (cm^2^); and **C**, maximum diameter (cm). P-values shown are for the regression of average methylation (DNAm) across the *ZNF300* differentially methylated region (DMR) onto each placental feature, adjusted for gestational age and feto-placental weight ratio. Males and females are denoted in blue and yellow, respectively.

### DNA methylation at *ZNF300* mediates genotype effects on placenta morphology

We searched for common genetic variants, specifically single nucleotide polymorphisms (SNPs), that are associated with DNA methylation levels at *ZNF300*, i.e. are methylation quantitative trait loci (meQTLs). After testing for *cis* (all SNPs on chromosome 5) and *trans* (all SNPs on the X chromosome) effects, we identified a total of 17 SNPs associated with DNA methylation at ZNF300 at a suggestive threshold of p ≤ 1E-4 (Supplementary Fig. 4), a threshold chosen based on the low statistical power for meQTL discovery in our placenta sample size (n = 120 with both DNAm and SNP data). Six of the meQTL SNPs were located on chromosome 5, including a peak in the ZNF300 differentially methylated region, and 11 meQTL SNPs were located on the X chromosome. Next, we tested associations between each of the 17 meQTL SNPs and placenta morphological features including area, perimeter, and max diameter. One *cis* meQTL SNP and 3 *trans* meQTL SNPs on the X-chromosome glycerol kinase gene (*GK*), showed associations with placenta perimeter (Supplementary Fig. 5) and max diameter (Supplementary Fig. 6). All 7 of the X chromosome meQTL SNPs and 1 chromosome 5 meQTL SNP were associated with placenta area (Supplementary Fig. 7). Given the links between genotype and placenta morphology, genotype and DNA methylation, and DNA methylation and placenta morphology, we tested the hypothesis that DNA methylation at *ZNF300* mediates the associations between genotype and placenta morphology. Causal inference testing revealed DNA methylation at the *ZNF300* locus mediates the effects of SNP rs13187817, located on outside of the *ZNF300* gene locus on chromosome 5, on placenta perimeter, area, and max diameter (Supplementary Fig. 5-7) and also mediates the effects of chromosome X SNPs on placenta area (Supplementary Fig. 7). An illustrative example is shown in Fig. 5 where an increase in DNA methylation, on average across the *ZNF300* region, is significantly associated with an increase in placenta area (*p*=0.014; Fig. 5a) and also minor allele count at chromosome X, physical position 30666654 ((*p*=6.176e^-05^; Fig. 5b). We also observed an increase in the number of minor alleles at the same locus significantly associated with placenta area (*p*=0.025; Fig. 5c). The genetic variant effect is attenuated and no longer significant when adjusted for DNA methylation at the *ZNF300* locus (*p*=0.093; Fig. 5d).

**Figure 5:**
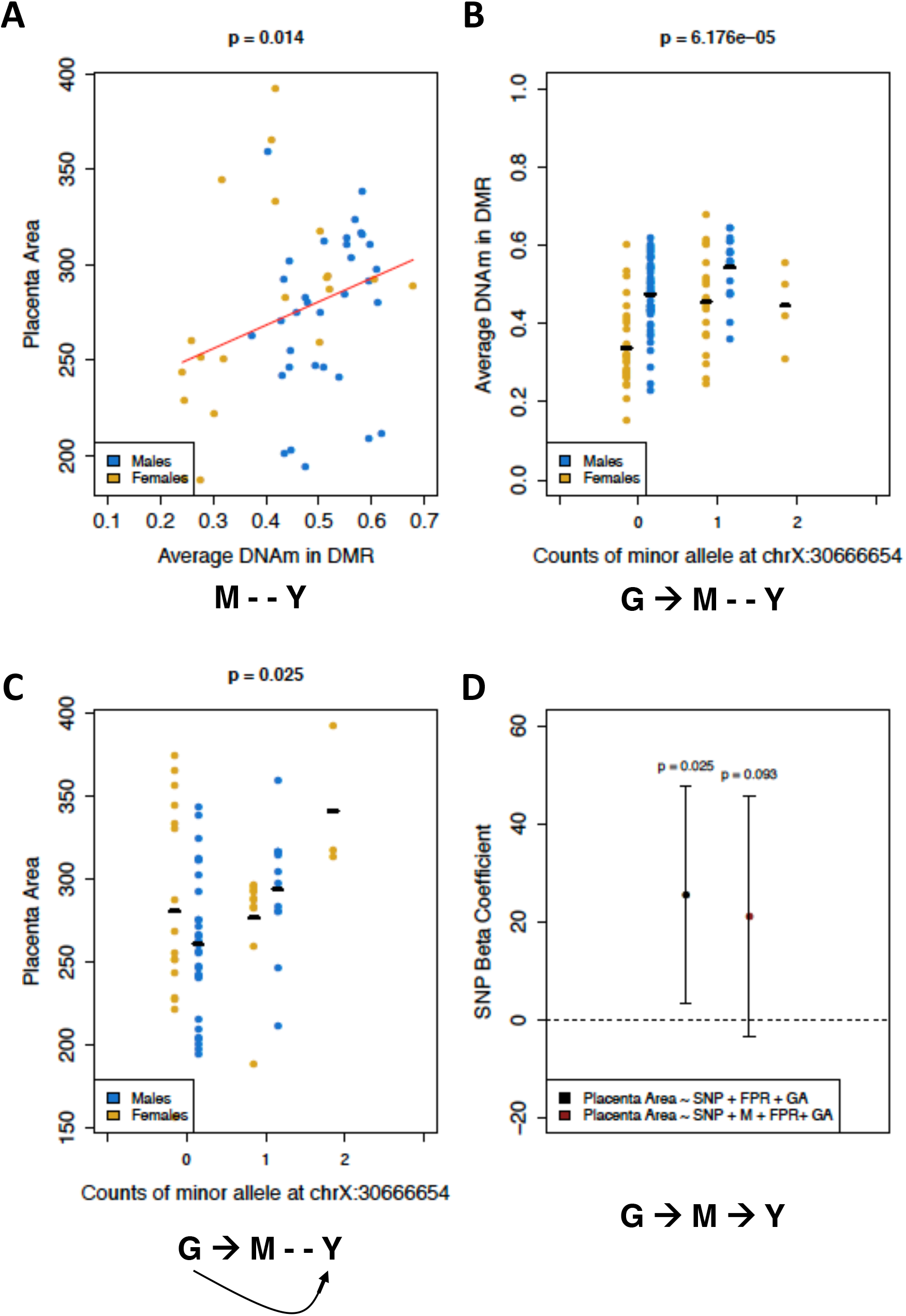
Placenta DNA methylation, at the *ZNF300* locus, mediates the association between locus-specific genetic variation and placenta area. Placenta from male and female offspring are denoted in blue and yellow, respectively. P-values shown from models that included feto-placental weight ratio and gestational age as covariates. **A**, Association between mean DNA methylation levels (y-axis) and placenta area (cm^2^). **B**, Causal association between a genetic variant at chromosome X, physical position 30666654, and average methylation across the *ZNF300* DMR. **C**, Causal association between chromosome X, physical position 30666654, genetic variant and placenta area. **D**, Genetic variant effect on placenta area with (black) and without (brown) adjustment for *ZNF300* DNA methylation in a linear regression model. Points denote beta coefficients with corresponding error bars and p-values also shown. The dashed line at zero represents a null finding, i.e. no effec t.

## Discussion

We identified a robust differentially methylated region (DMR) in the *ZNF300* promoter region associated with fetal sex in placenta tissue. The region showed a relatively large difference in methylation, with male offspring placentae exhibiting about a 15% increase in methylation relative to females. The sex associated DMR replicated across 6 additional, independent, studies that spanned measurement platforms, gestational ages, health outcomes, and included a data set with sorted villous cytotrophoblasts cells. Differences by sex at this locus were not observed in other developmentally relevant tissues including fetal brain, cord blood, or peripheral blood, suggesting the result is placenta specific. DNA methylation levels at this DMR are also associated with placenta area, perimeter, and max diameter and causal inference testing suggests DNA methylation at ZNF300 mediates both local and more distant genetic effects on placenta morphology.

*ZNF300* functions as a transcriptional repressor, and hypermethylation in the promoter of this gene has been previously associated with decreased gene expression [70]. Expression of the *ZNF300* has been previously associated with tumorigenesis in HeLa cells [71], but knockdown of *ZNF300* has been previously associated with cell proliferation in K562 (chronic myelogenous leukemia) cells[72]. Our results indicated that female samples with decreased *ZNF300* promoter methylation (i.e. those with increased *ZNF300* expression) tended to have placentas characterized by smaller areas and perimeters, indicating a decreased proliferative capacity. Therefore, placenta cells (chiefly whichever cell types drive this methylation association) seem to mimic K562 cells in their functionality, but future work is needed to confirm this theory.

DNA methylation in the *ZNF300* promoter region has been previously linked to intrauterine growth restriction (IUGR) in monochorionic twin samples [67]. We examined small for gestational age (SGA), which is related to IUGR, but did not observe associations between methylation and SGA in EARLI samples or a publicly available dataset (PR 3). Although not significant overall, when limiting analyses to females only, we did observe higher methylation levels at *ZNF300* among SGA females which is consistent in direction with previous IUGR findings [67]. We also found *lower* DNAm levels in this region to be associated with smaller placenta areas which is seemingly contradictory to previous reports that small placenta size is associated with IUGR, at least in women with low pregnancy-associated plasma protein-A[73]. Given the small number of SGA samples in our study, differences between SGA and IUGR phenotypes, and inconsistent findings with previous studies, future studies to rigorously investigate the relationship between methylation at *ZNF300* and fetal growth are warranted. Interestingly, females showed greater variability in DNA methylation (DNAm) at the *ZNF300* locus compared to males. Some placenta samples from female offspring showed lower DNAm compared to males. The remaining female samples had similar DNAm levels as the male placentae. It has been theorized that that males aim to maintain growth in the presence of an initial adverse insult, leaving less reserve capacity to handle an additional adverse event. In contrast, females reduce growth in the presence of a poor intrauterine environment, allowing for increased plasticity or robustness to perturbations or adverse conditions during pregnancy[74-76]. The greater distribution of placenta *ZNF300* methylation levels and their linked morphological phenotypes is congruent with this notion of female adaptability.

Two lines of evidence suggest the sex difference in methylation at the *ZNF300* locus is specific to placental cells. First, we observed no association between sex and *ZNF300* DNA methylation in cord blood, peripheral blood, and fetal tissues. Second, the vast majority of previous studies of autosomal DNAm differences related to sex in other tissues, including peripheral blood[50, 52, 77], cord blood[53], pancreas[54], and brain[55], did not report *ZNF300* differences. One study in peripheral blood did report 5 significant *ZNF300* sites in their initial discovery phase, but only replicated the result in 1 of their 3 replication cohorts, and only for 2 sites [51].

This study included comprehensive genome-wide coverage of methylation at over 26 million loci in a reasonably large discovery sample size. Increased sample sizes in future studies with high genome coverage could elucidate additional loci and differentially methylated regions that we were not sufficiently powered to detect. Given the observed increase in methylation variability among females, screening for differential variability related to fetal sex is also warranted in future studies. We also found suggestive evidence for local and more distant genetic effects on *ZNF300* DNAm levels; these findings should be confirmed in greater sample sizes of joint genotype and DNAm data than were available here (N = 121). Finally, future work is needed to examine the role of DNA methylation at this locus across the gestational period coupled with functional experiments to resolve the downstream consequences of differences in DNAm at *ZNF300* in placenta. Such studies can help determine the mechanisms by which *ZNF300* contributes to sex-specific *in utero* environments. A complementary article by Andrews et al.[78], which takes a mega-analysis approach, expanding the set of publicly available data that used Illumina 450K array data, also identifies differential placenta methylation at *ZNF300* associated with fetal sex, and demonstrates its consistency across each trimester of pregnancy. This further replication provides strong evidence supporting sex-specific methylation at the *ZNF300* locus and calls for additional investigation of *ZNF300* methylation as a possible placenta molecular mechanism relevant to sex differences in health outcomes across the lifespan.

## Methods

### EARLI study description

The Early Autism Risk Longitudinal Investigation (EARLI) is a prospective, enriched-risk birth cohort that has been described in detail elsewhere [79]. The EARLI study was approved by Human Subjects Institutional Review Boards (IRBs) from each of the four study sites (Johns Hopkins University, Drexel University, University of California Davis, and Kaiser Permanente). Informed consent was obtained from all participating families. The 232 mothers with a subsequent child born through this study had births between November 2009 and March 2012. Placental biopsy samples were collected after delivery at each clinical lab site using Baby Tischler Punch Biopsy Forceps. Samples were stored at ambient temperature in RNAlater vials (Qiagen, Cat. No. 76154) and shipped same-day to the Johns Hopkins Biological Repository (JHBR) in Baltimore, Maryland, for storage at -190°C until nuclei acid processing. After biopsy, the placentas were then placed flat in a bag with 10% formalin at a volume roughly equivalent to the size of the placenta. These bags were then shipped to Placenta Analytics, LLC, where they underwent extensive morphological testing via 2D fetal surface imaging, as previously described[80].

Infants were followed with extensive neurodevelopmental phenotyping until age three, when a diagnosis of ASD could reliably be made. Children were classified as having autism spectrum disorder (ASD) if they met or exceeded the ASD cutoff of the Autism Diagnostic Observation Schedule (ADOS) [81]and met the Diagnostic and Statistical Manual of Mental Disorders, 4^th^ edition, Text Revision[82] criteria for Autistic Disorder or Pervasive Developmental Disorder – Not otherwise specified. Children were classified as having non-typical development (NTD) if they did not meet the criteria for ASD classification but had two or more Mullen Scales of Early Learning[83] subtests ≥ 1.5 SD below the mean and/or one or more Mullen subtests ≥ 2 SD below the mean and/or ADOS ≤ 3 points below the ASD cutoff. Children were classified as having typical development (TD) if they did not meet the criteria for ASD classification and had no more than one Mullen subtest ≥ 1.5 SD below the mean and no Mullen subtest ≥ 2 SD below the mean and ADOS > 3 points below the ASD cutoff.

### Whole genome bisulfite sequencing processing, alignment, and quality control

Genomic DNA (gDNA) from biopsy punches was extracted at JHBR using a QIAgen QIAsymphony automated workstation with the Blood 1000 protocol of the DSP DNA Midi kit (Cat. No. 937255) as per manufacturer’s instructions. 250ng gDNA was bisulfite treated using the Zymo EZ-96 DNA Methylation Kit (Cat. No. D5004), and sequencing libraries were made with the NEBNext Ultra DNA Library Prep Kit for Illumina. Library preparation was performed at the Center for Epigenetics and libraries were sequenced at ∼4x coverage at the High Throughput Sequencing Center at the Johns Hopkins University School of Medicine using the 3 Illumina HiSeq 2500 instruments. Initially 133 placenta samples were run on 16 flow cells; two samples failed due to a fluidics error and were re-run using a 17^th^ flow cell. Raw .fastq files were aligned the human reference genome (hg38) downloaded from Ensembl[84] via Bismark (v0.16.3)[85] using the default arguments for paired-end sequencing data. M-bias plots[86] were used to inform the trimming of the 5bp of both the first and second reads, and methylation status was called using the Bismark methylation extractor function. All subsequent analyses used R (version 3.4.2) and the R package *bsseq* (version 1.13.6)[87]. The bsseq function *read*.*bismark*() was used to read in Bismark output files (“cytosine report” format) and obtain DNAm measurements on a per-CpG site basis. A total of 29,091,077 CpG sites were measured with at least 1 read in 1 sample out of the total 133. Next methylation levels were smoothed using the BSmooth algorithm that has been previously described [86]. Briefly, this technique employs a local-likelihood smoothing algorithm to recapitulate higher coverage whole genome bisulfite sequencing data from lower coverage data, exploiting known local correlation structure in DNAm data. We used the *BSmooth*() function in the *bsseq* R package with the default arguments.

### Identification of differentially methylated region

To characterize normal placentas, we limited our initial discovery sample to 37 samples (N _males_ = 17, N_females_ = 20) with a classification of TD at 3 years of age. We removed CpGs with little or no coverage in a large proportion of samples to avoid detection of single probes as regions, as they are likely to be false positives [87]. Specifically, we limited DNA methylation (DNAm) data to sites at which at least 8 males and 10 females had ≥ 2 reads at each CpG site, and also removed sex chromosome CpGs, leaving a total of 26,157,032 CpG sites. Finally we used the *BSmooth*.*tstat*.*fix*() and *dmrFinder*() functions in *bsseq* with the default arguments to identify differentially methylated regions (DMRs). Statistical significance was assessed by creating 1000 null sets of methylation data via scrambling sample labels for fetal sex and re-running the DMR-finding algorithm; p-values were calculated as the number of null sets in which at least 1 DMR had a ≥ ‘areaStat’ value (sum of t-statistics) and width (in bp) than the candidate DMR.

### Placenta replication datasets

To replicate and examine the robustness of DNA methylation changes related to fetal sex, we examined additional placenta datasets from independent sets of samples and studies (Supplementary Table 5). First, we examined the DMR in the 50 EARLI WGBS samples classified as NTD (N _males_ = 26, N_females_ = 24), termed Placenta Replication 1 (PR 1). Next, we downloaded the manifest for the Illumina HumanMethylation450 BeadChip (450K), the primary platform for which publicly available placenta DNAm data are available, and found that ten probes on the 450K probes overlapped with the *ZNF300* DMR coordinates. Therefore, we first downloaded three full term, mixed cell placenta data sets (matching the type of data we had for EARLI) from the Gene Expression Omnibus (GEO) [88] (GSE75248, GSE71678, and GSE75196), and obtained a fourth data set of this kind[67] from the study authors; these were termed PRs 2-5 respectively. Next to investigate if this DMR was present earlier in the gestational period, we downloaded the preterm samples from an additional data set (GSE57767), which we termed PR 6. Finally, to preclude the possibility that this DMR was driven by cell type heterogeneity[89], we downloaded an additional 450K data set of a single placenta cell type, villous cytotrophoblasts (GSE93208), which we termed PR 7. For 450K datasets for which raw data were available (PRs 2-5 and 7), we implemented the following processing and QC steps: functional normalization [90], removal of samples with low overall intensity (median methylated or unmethylated signal < 11) or with a detection p-value > 0.01 in more than 1% of probes, and removal of probes described as ambiguously mapping [91], or with a detection p-value > 0.01 in more than 10% of samples. For PR 6, for which raw data were not available, we performed quantile normalization [92] and removed ambiguously mapping probes.

### Data from other developmentally relevant tissues

We also sought to discover the extent to which the genome-wide significant placenta differentially methylated region (DMR) was present in different tissue types; we examined 450K data from cord blood, fetal brain, and peripheral blood. The cord blood data (N = 223) were obtained from EARLI samples; processing and quality control has been previously described[90]. Normalized percent methylation values for fetal brain data (N = 179) were downloaded from GEO (GSE58885). Peripheral blood data from children, aged 3-5 years, (N = 970) were used from the Study to Explore Early Development; processing and quality control has also been previously described[93].

### Genotype processing and methylation quantitative trait loci query

Genotype processing and QC for EARLI autosomal SNPs has been previously described [90]. Imputed genotype data for chromosome 5 were available on 682,195 SNPS for 121 of the 133 samples on which WGBS data existed. We imposed a 5% MAF threshold and removed duplicate positions, which limited the number of SNPs to 451,693, and then lifted the data from hg19 to hg38 (to match genome build on which WGBS alignment was performed) using the liftOver software[94], yielding a final count of 451,680 SNPs on chromosome 5. For the X chromosome, we measured 108,563 SNPs using the Illumina Omni5 plus exome array, which included 105,121 SNPs from the non-pseudoautosomal (nonPAR) region and 3,442 SNPs from the pseudoautosomal (PAR) region. Imposing a 5% MAF threshold yielded a total of 47,399 SNPs (44,992 nonPAR and 2,407 PAR), and lifting genome builds left a final total of 47,265 SNPs. To perform the methylation quantitative trait loci (meQTL) queries, we regressed the mean methylation level in the *ZNF300* DMR onto each genotype that passed QC and filtering thresholds from chromosomes 5 and X. We used an additive model for genotypes and adjusted for the first four principal components of ancestry and gestational age. For the X chromosome SNPs, we conducted this same analysis in each sex separately (N _males_= 70, N_females_= 50) and then meta-analyzed those results together using the METAL software [95] under the default conditions.

### Fetal growth and placenta morphology

We defined small for gestational age (SGA) any sample that did not exceed the tenth percentile of birth weight for its gestational age and sex according to United States reference data [96]. EARLI placenta morphology measures have been detailed elsewhere[97].

### Morphology mediation analyses

We regressed mean *ZNF300* methylation levels onto each morphological feature, adjusting for gestational age and feto-placental weight ratio; we did this analysis first in all samples on which DNAm and morphology data were available (N = 53), and then repeated the analyses in males (N = 33) and females (N = 20) separately. Finally, to explore the possibility that the suggestive genotype effects of meQTLs influenced these morphology phenotypes through *ZNF300* DNAm, we employed a causal inference testing (CIT), or mediation analysis, approach as we previously implemented for rheumatoid arthritis and child allergy [98, 99]. Briefly, this method conducts a series of regressions to establish the genotypic effect on a phenotype that acts through methylation, exploiting the implicit nature of the SNP effect to precede all other effects temporally. These regressions are:

1. Regression of phenotype (Y) onto methylation (M): Y and M are associated but no causality established
2. Regression of methylation (M) onto genotype (G): G is causally associated with M
3. Regression of Y onto G: G is causally associated with Y
4. Regression of Y onto G, adjusting for M: Attenuation of beta coefficient indicates that the causal effect of G on Y is mediated by M

We conducted a series of CIT procedures using all SNPs from the chromosome 5 and chromosome X meQTL queries that passed a suggestive significance threshold of p ≤ 1E-4 (n = 17), the mean methylation level from the *ZNF300* DMR, and the 4 placenta morphology phenotypes. For each regression, we included the total number of samples which had measurements for the form of data included in that regression. For regression 1, this was 53 samples. For regression 2, this was 120 samples. For regressions 3 and 4, this was 48 samples. For all regressions involving X chromosome genotype data, we conducted analyses separately in males and females and then meta-analyzed the results as done for the meQTL query.

## List of Abbreviations

450K: Illumina HumanMethylation450 BeadChip, ADOS: Autism Diagnostic Observation Schedule, ASD: autism spectrum disorder, BLAT: Blast-like alignment tool, CIT: causal inference testing, CpG: cytosine (phosphate) guanine dinucleotide, C-section: caesarean section, DNAm: DNA methylation, DMR: Differentially methylated region, EARLI: Early Autism Risk Longitudinal Investigation, GEO: gene expression omnibus, IUGR: intrauterine growth restriction, MAF: minor allele frequency, MD: mean difference, meQTL: methylation quantitative trait loci, non-PAR: non-pseudoautosomal region, NTD: non-typically developing, PAR: pseudoautosomal region, PR: placenta replication, QC: quality control, SGA: small for gestational age, SNP: single nucleotide polymorphism, TD: typically developing, TFBS: transcription factor binding site, WGBS: whole genome bisulfite sequencing, ZNF300: zinc finger 300

## Declarations

### Ethics Approval and Consent to Participate

Informed consent was obtained from all families that participated in EARLI. Human Subjects Institutional Review Boards (IRBs) from each of the four EARLI study sites (Johns Hopkins University, Drexel University, University of California Davis, and Kaiser Permanente) approved the EARLI study.

## Consent for Publication

Not applicable

## Availability of Data and Material

The EARLI whole-genome bisulfite sequencing methylation data is available via the National Database for Autism Research (NDAR) portal under collection # TBN. The other publicly available datasets we utilized can be obtained from their original sources: Gene Expression Omnibus (GEO) accessions GSE75248, GSE71678, GSE75196, GSE57767, and GSE93208 and from the Roifman et al study authors [67].

## Code Availability

Scripts for analysis conducted in this study are available at https://github.com/sandrews5/PlacentaDNAm_FetalSex

## Authors’ Contributions

SVA, MDF and CL-A conceived and designed the study. SVA and KMB performed processing and QC of WGBS and genotype data, respectively. CMS obtained placenta morphology measures in EARLI and provided input on analyses of these phenotypes. CJN, MDF, LAC, IH-P led EARLI sample recruitment and participation. MDF and APF obtained funding for the DNA methylation measurements in EARLI and supervised their acquisition by JIF and RT. RY informed analytical approaches and data interpretation of placenta morphology phenotypes. SVA performed data analyses under the mentorship of CL-A, KDH, and MDF with additional input from KMB. SVA, CL-A, and MDF wrote the paper. All authors contributed to interpretation of results and edited and reviewed the manuscript.

## Acknowledgements

We would like to thank Rosanna Weksberg and Sanaa Choufani for providing us data from their IUGR study. We would also like to thank Dan Arking for his thoughtful comments on analysis methods and data interpretation.

