## Additional Information for "Placenta DNA methylation at *ZNF300* is associated with fetal sex and placental morphology"

**This file contains the following information:**

**Supplementary Tables:**

Supplementary Figure 3…………………….……………………………………...…page 14

Supplementary Figure 4……………………………………………...……………page 15

Supplementary Figure 5……………………………………………………………...page 16

Supplementary Figure 6…………………………………………………………...…page 17

Supplementary Figure 7……………………………………………………………...page 18

Supplementary Figure 8………………………………………………………..……page 19

**Supplementary Table 1.** Descriptive characteristics of the Early Autism Risk Longitudinal Investigation (EARLI) samples with placenta whole genome bisulfite sequencing measures

| | *Number (%)* | | |
|---|---|---|---|
| | **Males (n=76)** | **Females (n=57)** | **p-value** |
| ***Neurodevelopmental diagnosis at age 3*** | | | 0.004[a] |
| Typically Developing | 17 (0.22) | 20 (0.35) | |
| Non-Typically Developing | 26 (0.34) | 24 (0.42) | |
| Autism Spectrum Disorder | 18 (0.24) | 1 (0.02) | |
| Lost to follow up | 15 (0.20) | 12 (0.21) | |
| ***Race*** | | | 0.467[a] |
| White | 43 (0.57) | 26 (0.46) | |
| Black | 6 (0.08) | 4 (0.07) | |
| Asian | 12 (0.16) | 8 (0.14) | |
| Other/Missing | 15 (0.20) | 19 (0.33) | |
| ***Mode of delivery*** | | | 0.290[a] |
| Vaginal | 38 (0.50) | 30 (0.53) | |
| C-section | 26 (0.34) | 12 (0.21) | |
| Missing | 12 (0.16) | 15 (0.26) | |
| ***Gestational Age (weeks)*** | | | |
| Mean (range) | 39.28 (35.99 - 42.43) | 39.41 (33.86 - 41.57) | 0.604[b] |

[a] chi square test p-value

[b] t-test p-value

**Supplementary Table 2.** BLAT results for ZNF300 DMR region. Results include BLAT results for reference sequence in DMR region and for source sequence of 9 450K probes that overlap this region.

| QUERY | SCORE | START | END | QSIZE | IDENTITY | CHRO | STRAND | START | END | SPAN |
|---|---|---|---|---|---|---|---|---|---|---|
| **450k Probes Overlapping *ZNF300 DMR*** | | | | | | | | | | |
| QUERY | SCORE | START | END | QSIZE | IDENTITY | CHRO | STRAND | START | END | SPAN |
| cg19486756 | 50 | 1 | 50 | 50 | 100.00% | 5 | - | 150281143 | 150281192 | 50 |
| cg12346881 | 50 | 1 | 50 | 50 | 100.00% | 5 | - | 150283837 | 150283886 | 50 |
| cg04675542 | 50 | 1 | 50 | 50 | 100.00% | 5 | + | 150284416 | 150284465 | 50 |
| cg02343823 | 50 | 1 | 50 | 50 | 100.00% | 5 | - | 150284419 | 150284468 | 50 |
| cg08580836 | 50 | 1 | 50 | 50 | 100.00% | 5 | - | 150284448 | 150284497 | 50 |
| cg08580836 | 32 | 12 | 46 | 50 | 97.10% | 6 | + | 1086171 | 1200162 | 113992 |
| cg19014419 | 50 | 1 | 50 | 50 | 100.00% | 5 | + | 150284456 | 150284505 | 50 |
| cg11291313 | 50 | 1 | 50 | 50 | 100.00% | 5 | - | 150284482 | 150284531 | 50 |
| cg18237551 | 50 | 1 | 50 | 50 | 100.00% | 5 | - | 150284552 | 150284601 | 50 |
| cg21228005 | 50 | 1 | 50 | 50 | 100.00% | 5 | + | 150284748 | 150284797 | 50 |
| **chr5:150904274-150905870 under hg38** | | | | | | | | | | |
| QUERY | SCORE | START | END | QSIZE | IDENTITY | CHRO | STRAND | START | END | SPAN |
| hg38_dna | 1597 | 1 | 1597 | 1597 | 100.00% | 5 | + | 150904274 | 150905870 | 1597 |
| hg38_dna | 269 | 463 | 869 | 1597 | 83.50% | 5 | + | 150946310 | 150946712 | 403 |
| hg38_dna | 30 | 1170 | 1254 | 1597 | 96.90% | 1 | + | 154418956 | 154419188 | 233 |
| hg38_dna | 28 | 1199 | 1244 | 1597 | 66.70% | 1 | + | 50659355 | 50659384 | 30 |
| hg38_dna | 26 | 1352 | 1381 | 1597 | 96.50% | 10 | + | 461616 | 461660 | 45 |
| hg38_dna | 26 | 1352 | 1381 | 1597 | 96.50% | 10 | + | 746531 | 746575 | 45 |
| hg38_dna | 26 | 810 | 836 | 1597 | 100.00% | 1 | + | 193931671 | 193932153 | 483 |
| hg38_dna | 23 | 1182 | 1204 | 1597 | 100.00% | 11 | - | 12070379 | 12070401 | 23 |
| hg38_dna | 23 | 1510 | 1533 | 1597 | 100.00% | 1 | + | 176074721 | 176074745 | 25 |
| hg38_dna | 22 | 405 | 426 | 1597 | 100.00% | 7 | - | 37414236 | 37414257 | 22 |
| hg38_dna | 22 | 1182 | 1203 | 1597 | 100.00% | 12 | - | 100354201 | 100354222 | 22 |
| hg38_dna | 22 | 244 | 265 | 1597 | 100.00% | 1 | - | 22907061 | 22907082 | 22 |
| hg38_dna | 21 | 182 | 202 | 1597 | 100.00% | 12 | - | 70460496 | 70460516 | 21 |
| hg38_dna | 20 | 1182 | 1203 | 1597 | 95.50% | 1 | - | 29551036 | 29551057 | 22 |

**Supplementary Table 3.** Association metrics for mean ZNF300 methylation levels and potential key confounders.

| Statistical model and confounder name | *Coefficient (95% Confidence Interval)* | *p-value*[a] |
|---|---|---|
| *Gestational Age:* | | |
| $E(M_{DMRmean}) = \alpha + \beta_1 GestationalAge + \varepsilon$ | 0.0025 (-0.0260, 0.0309) | 0.861 |
| *Genetic ancestry:* | | |
| $E(M_{DMRmean}) = \alpha + \beta_2 PC1 + \beta_2 PC2 + \beta_3 PC3 + \beta_4 PC4 + \beta_5 PC5 + \varepsilon$ | | |
| PC1 | -1.52 (-7.95, 4.91) | 0.631 |
| PC2 | -2.09 (-9.05, 4.87) | 0.541 |
| PC3 | 1.29 (-12.34, 14.93) | 0.846 |
| PC4 | -14.80 (-71.92, 42.31) | 0.598 |
| PC5 | -2.73 (-13.71, 8.26) | 0.613 |
| *Mode of delivery*: | | |
| $E(M_{DMRmean}) = \alpha + \beta_1 ModeofDelivery + \varepsilon$ | -0.032 (-0.165, 0.100) | 0.619 |

PC: principal component

[a]from linear regression of mean across 45 CpG sites in ZNF300 DMR vs gestational age, the first 5 principal components from genotype data, and mode of delivery (vaginal vs C-section).

**Supplementary Table 4.** Candidate differentially methylated regions associated with fetal sex identified in our genome-wide screen of placentae from neurotypical children.

| Chr | Start | End | Size (bp) | Mean_Males | Mean_Females | MeanDiff | fwer |
|---|---|---|---|---|---|---|---|
| 5 | 150904274 | 150905870 | 1597 | 0.53969502 | 0.389863571 | 0.14983145 | 0.015 |
| 6 | 102904326 | 102905438 | 1113 | 0.36358626 | 0.480718729 | -0.1171325 | 0.192 |
| 1 | 26817250 | 26817668 | 419 | 0.46238026 | 0.721890005 | -0.2595097 | 0.319 |
| 8 | 3182959 | 3183686 | 728 | 0.29756636 | 0.394339137 | -0.0967728 | 0.359 |
| 13 | 89087754 | 89088647 | 894 | 0.39849116 | 0.510989959 | -0.1124988 | 0.371 |
| 22 | 33014021 | 33014861 | 841 | 0.1672423 | 0.314354614 | -0.1471123 | 0.397 |
| 21 | 21024602 | 21025439 | 838 | 0.41402048 | 0.475943936 | -0.0619235 | 0.454 |
| 5 | 62804346 | 62805039 | 694 | 0.56725629 | 0.412467734 | 0.15478855 | 0.504 |
| 7 | 47052929 | 47053330 | 402 | 0.25084429 | 0.405674256 | -0.15483 | 0.524 |
| 21 | 21019803 | 21020643 | 841 | 0.36591433 | 0.416062629 | -0.0501483 | 0.533 |
| 4 | 37001712 | 37002384 | 673 | 0.23213303 | 0.321256093 | -0.0891231 | 0.672 |
| 18 | 30175567 | 30176200 | 634 | 0.2225856 | 0.366395684 | -0.1438101 | 0.739 |
| 4 | 25506054 | 25506453 | 400 | 0.29781156 | 0.475505546 | -0.177694 | 0.746 |
| 11 | 34346080 | 34346768 | 689 | 0.74035313 | 0.625274156 | 0.11507897 | 0.803 |
| 4 | 56433717 | 56434325 | 609 | 0.62517709 | 0.481522305 | 0.14365478 | 0.823 |
| 9 | 16869185 | 16869742 | 558 | 0.44334902 | 0.286416548 | 0.15693247 | 0.87 |
| 17 | 30823009 | 30823535 | 527 | 0.58974505 | 0.438174901 | 0.15157015 | 0.891 |
| 9 | 63859185 | 63859620 | 436 | 0.18865911 | 0.291493489 | -0.1028344 | 0.904 |
| 3 | 33114982 | 33115497 | 516 | 0.59630619 | 0.441793945 | 0.15451224 | 0.918 |
| 7 | 25743955 | 25744332 | 378 | 0.22377351 | 0.351845682 | -0.1280722 | 0.931 |
| 3 | 196163915 | 196164420 | 506 | 0.22882054 | 0.326757973 | -0.0979374 | 0.946 |
| 8 | 4313287 | 4313811 | 525 | 0.40328952 | 0.450709515 | -0.04742 | 0.948 |
| 2 | 60883035 | 60883593 | 559 | 0.40270084 | 0.240521326 | 0.16217951 | 0.955 |
| 7 | 43113076 | 43113374 | 299 | 0.35889834 | 0.213780459 | 0.14511788 | 0.956 |
| 9 | 5683764 | 5684207 | 444 | 0.66013324 | 0.794337795 | -0.1342046 | 0.961 |
| 17 | 82228306 | 82228514 | 209 | 0.56440784 | 0.404294969 | 0.16011288 | 0.964 |
| 17 | 503023 | 503261 | 239 | 0.31963847 | 0.468408437 | -0.14877 | 0.972 |
| 17 | 80832988 | 80833421 | 434 | 0.49849612 | 0.376316696 | 0.12217943 | 0.973 |
| 1 | 112957285 | 112957782 | 498 | 0.62968011 | 0.512139614 | 0.1175405 | 0.976 |
| 8 | 4643964 | 4644467 | 504 | 0.37603323 | 0.431062148 | -0.0550289 | 0.976 |
| 22 | 47631852 | 47632172 | 321 | 0.29063687 | 0.418916495 | -0.1282796 | 0.977 |
| 11 | 26114759 | 26115252 | 494 | 0.55515393 | 0.625585307 | -0.0704314 | 0.977 |
| 2 | 55442125 | 55442427 | 303 | 0.65478068 | 0.785600569 | -0.1308199 | 0.979 |
| 13 | 96947238 | 96947640 | 403 | 0.56990033 | 0.45208243 | 0.1178179 | 0.982 |
| 2 | 43218773 | 43219225 | 453 | 0.6417339 | 0.515743148 | 0.12599076 | 0.987 |
| 11 | 118788960 | 118789320 | 361 | 0.57662116 | 0.431160296 | 0.14546086 | 0.988 |
| 11 | 748309 | 748750 | 442 | 0.56287014 | 0.425001035 | 0.13786911 | 0.989 |
| 19 | 52536440 | 52536743 | 304 | 0.15673983 | 0.341667011 | -0.1849272 | 0.991 |

| | | | | | | | |
|---|---|---|---|---|---|---|---|
| 11 | 23730361 | 23730589 | 229 | 0.3203217 | 0.44812786 | -0.1278062 | 0.995 |
| 21 | 10119719 | 10119962 | 244 | 0.12107314 | 0.191750242 | -0.0706771 | 0.997 |
| 21 | 20997548 | 20997812 | 265 | 0.14630042 | 0.220026114 | -0.0737257 | 0.997 |
| 7 | 30596194 | 30596560 | 367 | 0.31803916 | 0.194787724 | 0.12325144 | 0.997 |
| 18 | 49017485 | 49017854 | 370 | 0.35070831 | 0.478526163 | -0.1278179 | 0.997 |
| 5 | 150906191 | 150906538 | 348 | 0.59324673 | 0.487077205 | 0.10616953 | 0.998 |
| 22 | 10961644 | 10961748 | 105 | 0.2034263 | 0.282579507 | -0.0791532 | 1 |
| 2 | 226835303 | 226835549 | 247 | 0.2530793 | 0.399235894 | -0.1461566 | 1 |
| 4 | 6953213 | 6953403 | 191 | 0.79051205 | 0.656509643 | 0.1340024 | 1 |
| 8 | 6523620 | 6523876 | 257 | 0.57330059 | 0.738626583 | -0.165326 | 1 |
| 12 | 68747362 | 68747557 | 196 | 0.47457651 | 0.358629776 | 0.11594673 | 1 |
| 11 | 95037869 | 95037965 | 97 | 0.15420386 | 0.27446568 | -0.1202618 | 1 |
| 7 | 24977573 | 24977748 | 176 | 0.77516372 | 0.626604073 | 0.14855964 | 1 |
| 6 | 48069444 | 48069679 | 236 | 0.4746624 | 0.345747952 | 0.12891444 | 1 |
| 1 | 26863968 | 26864266 | 299 | 0.61154535 | 0.446746529 | 0.16479882 | 1 |
| 22 | 21569064 | 21569310 | 247 | 0.52440884 | 0.385207195 | 0.13920165 | 1 |
| 9 | 63708818 | 63708951 | 134 | 0.12622869 | 0.237047741 | -0.1108191 | 1 |
| 5 | 39787881 | 39788013 | 133 | 0.5236956 | 0.656046628 | -0.132351 | 1 |
| 1 | 27552048 | 27552178 | 131 | 0.66701315 | 0.776846896 | -0.1098338 | 1 |
| 20 | 63641308 | 63641388 | 81 | 0.44121346 | 0.26799749 | 0.17321597 | 1 |
| 1 | 156436807 | 156436972 | 166 | 0.44166161 | 0.302576825 | 0.13908479 | 1 |
| 1 | 44794325 | 44794593 | 269 | 0.63957297 | 0.501941886 | 0.13763109 | 1 |
| 3 | 128490875 | 128491063 | 189 | 0.52224952 | 0.340747385 | 0.18150214 | 1 |
| 1 | 148844264 | 148844375 | 112 | 0.18319096 | 0.368005892 | -0.1848149 | 1 |
| 18 | 12078 | 12161 | 84 | 0.43858114 | 0.56079891 | -0.1222178 | 1 |
| 10 | 117373474 | 117373586 | 113 | 0.66364214 | 0.534971421 | 0.12867071 | 1 |
| 2 | 25216986 | 25217110 | 125 | 0.29588857 | 0.396137034 | -0.1002485 | 1 |
| 9 | 133819210 | 133819365 | 156 | 0.46475588 | 0.559329468 | -0.0945736 | 1 |
| 17 | 36658370 | 36658501 | 132 | 0.50973198 | 0.3657855 | 0.14394648 | 1 |
| 22 | 11618202 | 11618321 | 120 | 0.49343217 | 0.606624468 | -0.1131923 | 1 |
| 7 | 72880605 | 72880834 | 230 | 0.69341631 | 0.552380268 | 0.14103604 | 1 |
| 3 | 157542604 | 157542648 | 45 | 0.2755862 | 0.383824841 | -0.1082386 | 1 |
| 3 | 46845138 | 46845337 | 200 | 0.48711593 | 0.374539782 | 0.11257615 | 1 |
| 7 | 151460861 | 151461128 | 268 | 0.51639096 | 0.406896982 | 0.10949398 | 1 |
| 3 | 170909038 | 170909191 | 154 | 0.33495208 | 0.440143312 | -0.1051912 | 1 |
| 7 | 44846295 | 44846413 | 119 | 0.57697319 | 0.458717672 | 0.11825552 | 1 |
| 9 | 136110285 | 136110367 | 83 | 0.68297503 | 0.515186473 | 0.16778856 | 1 |
| 2 | 3528256 | 3528357 | 102 | 0.71419382 | 0.555728818 | 0.158465 | 1 |
| 3 | 150703021 | 150703172 | 152 | 0.62033306 | 0.498521837 | 0.12181123 | 1 |
| 3 | 195659311 | 195659409 | 99 | 0.67656181 | 0.547707109 | 0.1288547 | 1 |
| 9 | 42636362 | 42636446 | 85 | 0.36246595 | 0.499199983 | -0.136734 | 1 |

| | | | | | | | |
|---|---|---|---|---|---|---|---|
| 1 | 51877102 | 51877331 | 230 | 0.6328365 | 0.513548997 | 0.1192875 | 1 |
| 9 | 66269167 | 66269268 | 102 | 0.43975249 | 0.312889684 | 0.12686281 | 1 |
| 14 | 18966635 | 18966714 | 80 | 0.41976397 | 0.558251667 | -0.1384877 | 1 |
| 11 | 95037404 | 95037492 | 89 | 0.13404375 | 0.251162466 | -0.1171187 | 1 |
| 19 | 2428394 | 2428527 | 134 | 0.55021488 | 0.419253284 | 0.1309616 | 1 |
| 1 | 9183468 | 9183541 | 74 | 0.55327814 | 0.435611897 | 0.11766624 | 1 |
| 6 | 158939841 | 158939952 | 112 | 0.63060994 | 0.499820474 | 0.13078946 | 1 |
| 11 | 5937993 | 5938095 | 103 | 0.28469028 | 0.385370379 | -0.1006801 | 1 |
| 7 | 155205469 | 155205605 | 137 | 0.42057054 | 0.254319313 | 0.16625123 | 1 |
| 1 | 246794195 | 246794381 | 187 | 0.51090202 | 0.384268641 | 0.12663338 | 1 |
| 3 | 9648787 | 9648886 | 100 | 0.47914683 | 0.594951797 | -0.115805 | 1 |
| 7 | 91881387 | 91881554 | 168 | 0.29518391 | 0.426140411 | -0.1309565 | 1 |
| 1 | 3538797 | 3538882 | 86 | 0.47672223 | 0.558948672 | -0.0822264 | 1 |
| 16 | 69425919 | 69426137 | 219 | 0.77196256 | 0.624085947 | 0.14787661 | 1 |
| 14 | 59951529 | 59951675 | 147 | 0.30353118 | 0.482298425 | -0.1787672 | 1 |
| 1 | 66931569 | 66931885 | 317 | 0.39192873 | 0.261945555 | 0.12998318 | 1 |
| 2 | 74530953 | 74531109 | 157 | 0.53756719 | 0.401683879 | 0.13588331 | 1 |
| 3 | 24522364 | 24522452 | 89 | 0.50086357 | 0.385242479 | 0.11562109 | 1 |
| 6 | 27630425 | 27630482 | 58 | 0.4420674 | 0.580163934 | -0.1380965 | 1 |
| 16 | 68357089 | 68357286 | 198 | 0.69216437 | 0.789533419 | -0.097369 | 1 |
| 16 | 71896786 | 71896916 | 131 | 0.58803064 | 0.472353201 | 0.11567744 | 1 |
| 20 | 38599875 | 38599894 | 20 | 0.46951431 | 0.587585335 | -0.118071 | 1 |
| 16 | 75624202 | 75624266 | 65 | 0.29593336 | 0.420665851 | -0.1247325 | 1 |
| 5 | 95960156 | 95960220 | 65 | 0.6281615 | 0.513862969 | 0.11429853 | 1 |
| 5 | 178087094 | 178087146 | 53 | 0.63132238 | 0.52194273 | 0.10937965 | 1 |
| 15 | 73221239 | 73221312 | 74 | 0.54046885 | 0.635315408 | -0.0948466 | 1 |
| 6 | 33453156 | 33453403 | 248 | 0.64653333 | 0.471606606 | 0.17492672 | 1 |
| 1 | 143331925 | 143332013 | 89 | 0.51384825 | 0.766218557 | -0.2523703 | 1 |
| 3 | 9916172 | 9916363 | 192 | 0.54078141 | 0.408062861 | 0.13271855 | 1 |
| 6 | 7313845 | 7313973 | 129 | 0.20787541 | 0.316457052 | -0.1085816 | 1 |
| 15 | 74907469 | 74907697 | 229 | 0.61951186 | 0.505559664 | 0.1139522 | 1 |
| 21 | 10274090 | 10274234 | 145 | 0.62722703 | 0.424985944 | 0.20224108 | 1 |
| 7 | 44574706 | 44574745 | 40 | 0.41166986 | 0.530819084 | -0.1191492 | 1 |
| 1 | 32499813 | 32499843 | 31 | 0.47055569 | 0.573593885 | -0.1030382 | 1 |
| 1 | 91884479 | 91884576 | 98 | 0.54902255 | 0.425348841 | 0.12367371 | 1 |
| 4 | 37669799 | 37669900 | 102 | 0.57870869 | 0.425658962 | 0.15304973 | 1 |
| 15 | 55589346 | 55589409 | 64 | 0.40191328 | 0.286285847 | 0.11562744 | 1 |
| 14 | 59951010 | 59951115 | 106 | 0.40955544 | 0.610216582 | -0.2006611 | 1 |
| 10 | 131972756 | 131972817 | 62 | 0.73230174 | 0.617520246 | 0.11478149 | 1 |
| 16 | 77527 | 77580 | 54 | 0.41594531 | 0.292566949 | 0.12337836 | 1 |
| 1 | 52663482 | 52663498 | 17 | 0.66321365 | 0.556284149 | 0.1069295 | 1 |

| | | | | | | | |
|---|---|---|---|---|---|---|---|
| 16 | 33516339 | 33516408 | 70 | 0.21562682 | 0.327995265 | -0.1123684 | 1 |
| 1 | 144556793 | 144557065 | 273 | 0.50672015 | 0.663920838 | -0.1572007 | 1 |
| 1 | 246793605 | 246793764 | 160 | 0.57924049 | 0.451079821 | 0.12816067 | 1 |
| 6 | 31701851 | 31702125 | 275 | 0.46163439 | 0.351559334 | 0.11007505 | 1 |
| 19 | 39406116 | 39406142 | 27 | 0.54559645 | 0.397974967 | 0.14762149 | 1 |
| 3 | 25662744 | 25662866 | 123 | 0.45914893 | 0.344636435 | 0.11451249 | 1 |
| 4 | 17811924 | 17812185 | 262 | 0.64039078 | 0.510493737 | 0.12989705 | 1 |
| 2 | 233867831 | 233867859 | 29 | 0.38082591 | 0.497871826 | -0.1170459 | 1 |
| 6 | 116370081 | 116370221 | 141 | 0.60289399 | 0.487620281 | 0.11527371 | 1 |
| 19 | 54188248 | 54188433 | 186 | 0.55829196 | 0.427362155 | 0.1309298 | 1 |
| 1 | 54490242 | 54490348 | 107 | 0.56234151 | 0.431485684 | 0.13085582 | 1 |
| 5 | 175968949 | 175969073 | 125 | 0.69677009 | 0.574684422 | 0.12208567 | 1 |
| 1 | 148150643 | 148150756 | 114 | 0.50371152 | 0.389486313 | 0.11422521 | 1 |
| 3 | 61637971 | 61638111 | 141 | 0.67011876 | 0.54714846 | 0.1229703 | 1 |
| 11 | 65541703 | 65541746 | 44 | 0.23444536 | 0.336844875 | -0.1023995 | 1 |
| 1 | 40980935 | 40980998 | 64 | 0.64091945 | 0.525332982 | 0.11558647 | 1 |
| 19 | 33195448 | 33195484 | 37 | 0.4894217 | 0.361151367 | 0.12827033 | 1 |
| 7 | 3914236 | 3914329 | 94 | 0.40727373 | 0.494978403 | -0.0877047 | 1 |
| 1 | 17683690 | 17683702 | 13 | 0.48889576 | 0.561590693 | -0.0726949 | 1 |
| 4 | 109561771 | 109561817 | 47 | 0.42410893 | 0.304901868 | 0.11920707 | 1 |
| 21 | 5156545 | 5156548 | 4 | 0.96610397 | 0.765822717 | 0.20028126 | 1 |
| 5 | 177517878 | 177517889 | 12 | 0.64105401 | 0.509999735 | 0.13105427 | 1 |
| 12 | 47904075 | 47904079 | 5 | 0.46911267 | 0.338785423 | 0.13032725 | 1 |
| 17 | 26593455 | 26593488 | 34 | 0.46786941 | 0.593773318 | -0.1259039 | 1 |
| 21 | 21019221 | 21019408 | 188 | 0.3442985 | 0.376214985 | -0.0319165 | 1 |
| 1 | 148246176 | 148246207 | 32 | 0.57935934 | 0.718871882 | -0.1395125 | 1 |
| 6 | 27632298 | 27632400 | 103 | 0.66129806 | 0.552199735 | 0.10909833 | 1 |
| 19 | 43625504 | 43625562 | 59 | 0.26881748 | 0.408809886 | -0.1399924 | 1 |
| 10 | 46378876 | 46378915 | 40 | 0.58147216 | 0.445863148 | 0.13560901 | 1 |
| 14 | 59950659 | 59950706 | 48 | 0.49916485 | 0.657172098 | -0.1580072 | 1 |
| 5 | 178357743 | 178357761 | 19 | 0.4577331 | 0.326923447 | 0.13080966 | 1 |
| 17 | 83052683 | 83052706 | 24 | 0.69941798 | 0.574889102 | 0.12452887 | 1 |
| 10 | 80356201 | 80356313 | 113 | 0.49187432 | 0.377029589 | 0.11484473 | 1 |
| 1 | 93511123 | 93511212 | 90 | 0.41047113 | 0.51996816 | -0.109497 | 1 |
| 1 | 35030799 | 35030804 | 6 | 0.59654116 | 0.488844218 | 0.10769694 | 1 |
| 1 | 26827742 | 26827791 | 50 | 0.52751101 | 0.416573137 | 0.11093787 | 1 |
| 12 | 51024867 | 51024885 | 19 | 0.42176187 | 0.297616293 | 0.12414558 | 1 |
| 1 | 224214105 | 224214193 | 89 | 0.55652502 | 0.438119016 | 0.11840601 | 1 |
| 19 | 48752175 | 48752198 | 24 | 0.46926756 | 0.340727212 | 0.12854035 | 1 |
| 6 | 73452503 | 73452508 | 6 | 0.3472082 | 0.248373512 | 0.09883469 | 1 |
| 3 | 172038972 | 172038989 | 18 | 0.46459088 | 0.563677592 | -0.0990867 | 1 |

| | | | | | | | |
|---|---|---|---|---|---|---|---|
| 10 | 14579108 | 14579232 | 125 | 0.47160821 | 0.346163131 | 0.12544508 | 1 |
| 9 | 129738255 | 129738264 | 10 | 0.71937816 | 0.577432802 | 0.14194536 | 1 |
| 16 | 14909925 | 14909925 | 1 | 0.76295957 | 0.493747879 | 0.26921169 | 1 |
| 5 | 176239941 | 176239941 | 1 | 0.40962175 | 0.609286272 | -0.1996645 | 1 |
| 3 | 141488485 | 141488485 | 1 | 0.48767862 | 0.343482291 | 0.14419633 | 1 |
| 14 | 64896891 | 64896891 | 1 | 0.60252301 | 0.762276118 | -0.1597531 | 1 |
| 7 | 155345599 | 155345599 | 1 | 0.58876891 | 0.421927371 | 0.16684154 | 1 |
| 2 | 42013609 | 42013609 | 1 | 0.62957903 | 0.762344137 | -0.1327651 | 1 |
| 7 | 102573673 | 102573673 | 1 | 0.52415798 | 0.351503911 | 0.17265407 | 1 |
| 14 | 56669487 | 56669487 | 1 | 0.71694676 | 0.557403479 | 0.15954328 | 1 |
| 22 | 42555032 | 42555032 | 1 | 0.48484988 | 0.623788462 | -0.1389386 | 1 |
| 7 | 73157667 | 73157667 | 1 | 0.48097514 | 0.306001796 | 0.17497335 | 1 |
| 12 | 48106738 | 48106738 | 1 | 0.28632274 | 0.423368207 | -0.1370455 | 1 |
| 9 | 41360138 | 41360138 | 1 | 0.60075944 | 0.742406304 | -0.1416469 | 1 |
| 22 | 10571277 | 10571277 | 1 | 0.07631221 | 0.226366377 | -0.1500542 | 1 |
| 4 | 138562248 | 138562248 | 1 | 0.58136789 | 0.700456576 | -0.1190887 | 1 |
| 7 | 44122997 | 44122997 | 1 | 0.46375832 | 0.34881195 | 0.11494637 | 1 |
| 14 | 56668599 | 56668599 | 1 | 0.69761312 | 0.545062483 | 0.15255064 | 1 |
| 15 | 31390805 | 31390805 | 1 | 0.6136945 | 0.725216419 | -0.1115219 | 1 |
| 5 | 141479035 | 141479035 | 1 | 0.34474105 | 0.218042163 | 0.12669888 | 1 |
| 8 | 132774577 | 132774577 | 1 | 0.57824679 | 0.694545031 | -0.1162982 | 1 |
| 1 | 144811640 | 144811640 | 1 | 0.81487167 | 0.611641586 | 0.20323008 | 1 |
| 20 | 3172769 | 3172769 | 1 | 0.56526936 | 0.417757099 | 0.14751226 | 1 |
| 2 | 60884087 | 60884087 | 1 | 0.71727167 | 0.591259503 | 0.12601217 | 1 |
| 8 | 60516096 | 60516096 | 1 | 0.69144685 | 0.568598516 | 0.12284833 | 1 |
| 6 | 125987445 | 125987445 | 1 | 0.63931903 | 0.528588084 | 0.11073095 | 1 |
| 16 | 22161028 | 22161028 | 1 | 0.55505067 | 0.668965798 | -0.1139151 | 1 |
| 16 | 67561316 | 67561316 | 1 | 0.75570437 | 0.630246893 | 0.12545748 | 1 |
| 15 | 32535114 | 32535114 | 1 | 0.74831672 | 0.607058895 | 0.14125783 | 1 |
| 10 | 47301071 | 47301071 | 1 | 0.46625522 | 0.337512243 | 0.12874298 | 1 |
| 3 | 185253475 | 185253475 | 1 | 0.34076158 | 0.228240374 | 0.1125212 | 1 |
| 1 | 149814128 | 149814128 | 1 | 0.66076446 | 0.555724691 | 0.10503977 | 1 |
| 9 | 35829998 | 35829998 | 1 | 0.65957111 | 0.544194128 | 0.11537698 | 1 |
| 9 | 26945818 | 26945818 | 1 | 0.41827689 | 0.294387977 | 0.12388891 | 1 |
| 2 | 91627174 | 91627174 | 1 | 0.40569202 | 0.510175384 | -0.1044834 | 1 |
| 1 | 149102458 | 149102458 | 1 | 0.2077392 | 0.095397407 | 0.1123418 | 1 |
| 6 | 69866138 | 69866138 | 1 | 0.26873121 | 0.376107086 | -0.1073759 | 1 |
| 9 | 4986535 | 4986535 | 1 | 0.55584205 | 0.447691365 | 0.10815069 | 1 |
| 11 | 3839901 | 3839901 | 1 | 0.61323566 | 0.46494706 | 0.1482886 | 1 |
| 17 | 42387217 | 42387217 | 1 | 0.36227666 | 0.248823696 | 0.11345297 | 1 |
| 6 | 691690 | 691690 | 1 | 0.7512944 | 0.650367472 | 0.10092693 | 1 |

| | | | | | | | |
|---|---|---|---|---|---|---|---|
| 6 | 128521514 | 128521514 | 1 | 0.62356283 | 0.519314497 | 0.10424833 | 1 |
| 22 | 50807836 | 50807836 | 1 | 0.16221806 | 0.149821874 | 0.01239618 | 1 |
| 2 | 126181612 | 126181612 | 1 | 0.67824842 | 0.549772994 | 0.12847542 | 1 |
| 3 | 128075246 | 128075246 | 1 | 0.32231602 | 0.218337287 | 0.10397873 | 1 |
| 1 | 40664754 | 40664754 | 1 | 0.50203857 | 0.622019586 | -0.119981 | 1 |
| 6 | 2952096 | 2952096 | 1 | 0.7660068 | 0.650901061 | 0.11510574 | 1 |
| 15 | 65516307 | 65516307 | 1 | 0.57163284 | 0.467732622 | 0.10390022 | 1 |
| 2 | 169325251 | 169325251 | 1 | 0.42197327 | 0.522301529 | -0.1003283 | 1 |
| 2 | 42013064 | 42013064 | 1 | 0.68388401 | 0.787728952 | -0.1038449 | 1 |
| 21 | 44013979 | 44013979 | 1 | 0.63639722 | 0.513152493 | 0.12324472 | 1 |
| 20 | 38462914 | 38462914 | 1 | 0.2362895 | 0.413050671 | -0.1767612 | 1 |
| 2 | 42974439 | 42974439 | 1 | 0.67467076 | 0.560916823 | 0.11375394 | 1 |
| 14 | 70254233 | 70254233 | 1 | 0.66226424 | 0.765156671 | -0.1028924 | 1 |
| 17 | 51154571 | 51154571 | 1 | 0.55139713 | 0.444112232 | 0.1072849 | 1 |
| 6 | 27471878 | 27471878 | 1 | 0.40566709 | 0.305311989 | 0.1003551 | 1 |
| 19 | 3762180 | 3762180 | 1 | 0.45244129 | 0.327413649 | 0.12502764 | 1 |
| 1 | 235865757 | 235865757 | 1 | 0.45189121 | 0.353956541 | 0.09793467 | 1 |
| 3 | 134794466 | 134794466 | 1 | 0.19410446 | 0.266266354 | -0.0721619 | 1 |
| 3 | 136750396 | 136750396 | 1 | 0.52884958 | 0.427662684 | 0.1011869 | 1 |
| 2 | 102737500 | 102737500 | 1 | 0.39819639 | 0.2828347 | 0.11536169 | 1 |
| 11 | 100481801 | 100481801 | 1 | 0.57406559 | 0.652814729 | -0.0787491 | 1 |
| 6 | 79948856 | 79948856 | 1 | 0.34863754 | 0.248128244 | 0.1005093 | 1 |
| 16 | 67659354 | 67659354 | 1 | 0.72673198 | 0.615644515 | 0.11108746 | 1 |
| 20 | 25984970 | 25984970 | 1 | 0.22554291 | 0.349131263 | -0.1235884 | 1 |
| 13 | 89162841 | 89162841 | 1 | 0.27718697 | 0.369332381 | -0.0921454 | 1 |
| 1 | 29479899 | 29479899 | 1 | 0.3808028 | 0.48124185 | -0.1004391 | 1 |
| 16 | 33516325 | 33516325 | 1 | 0.2331225 | 0.343133687 | -0.1100112 | 1 |
| 16 | 33516312 | 33516312 | 1 | 0.23897963 | 0.34769418 | -0.1087145 | 1 |
| 13 | 109141156 | 109141156 | 1 | 0.51847749 | 0.662891152 | -0.1444137 | 1 |
| 1 | 152037550 | 152037550 | 1 | 0.65057056 | 0.545019567 | 0.10555099 | 1 |
| 13 | 89162879 | 89162879 | 1 | 0.27518599 | 0.366168118 | -0.0909821 | 1 |
| 3 | 129427888 | 129427888 | 1 | 0.57537001 | 0.467229106 | 0.1081409 | 1 |

FWER: family wise error rate

[a]Mean difference value (‘MeanDiff’) calculated as (males –females methylation) across all measured positions in DMR region.

**Supplementary Table 5.** Summary of all data sets (discovery and replication) used in replication analyses.

| Dataset Name | Tissue Type | Measurement Platform | Cohort/GEO ID | $N_{total}$ | $N_{Males}$ | $N_{Females}$ |
|---|---|---|---|---|---|---|
| ***Discovery:*** | | | | | | |
| *EARLI* | Placenta | WGBS | EARLI | 37 | 17 | 20 |
| ***Replication:*** | | | | | | |
| *PR 1* | Placenta | WGBS | EARLI | 50 | 26 | 24 |
| *PR 2* | Placenta | 450K | GSE75248 | 324 | 159 | 165 |
| *PR 3* | Placenta | 450K | GSE71678 | 342 | 184 | 158 |
| *PR 4* | Placenta | 450K | GSE75196 | 24 | 11 | 13 |
| *PR 5* | Placenta | 450K | Roifman et al. 2016 | 16 | 10 | 6 |
| *PR 6* | Placenta | 450K | GSE57767 | 12 | 6 | 6 |
| *PR 7* | Placenta | 450K | GSE93208 | 19 | 9 | 10 |
| ***Specificity Tissue 1*** | Cord Blood | 450K | EARLI | 223 | 114 | 109 |
| ***Specificity Tissue 2*** | Fetal Brain | 450K | GSE58885 | 179 | 100 | 79 |
| ***Specificity Tissue 3*** | Peripheral blood | 450K | SEED | 970 | 643 | 327 |

EARLI, Early Autism Risk Longitudinal Investigation; PR, placental replication; WGBS, whole genome bisulfite sequencing; 450K, Illumina 450K Methylation BeadChip

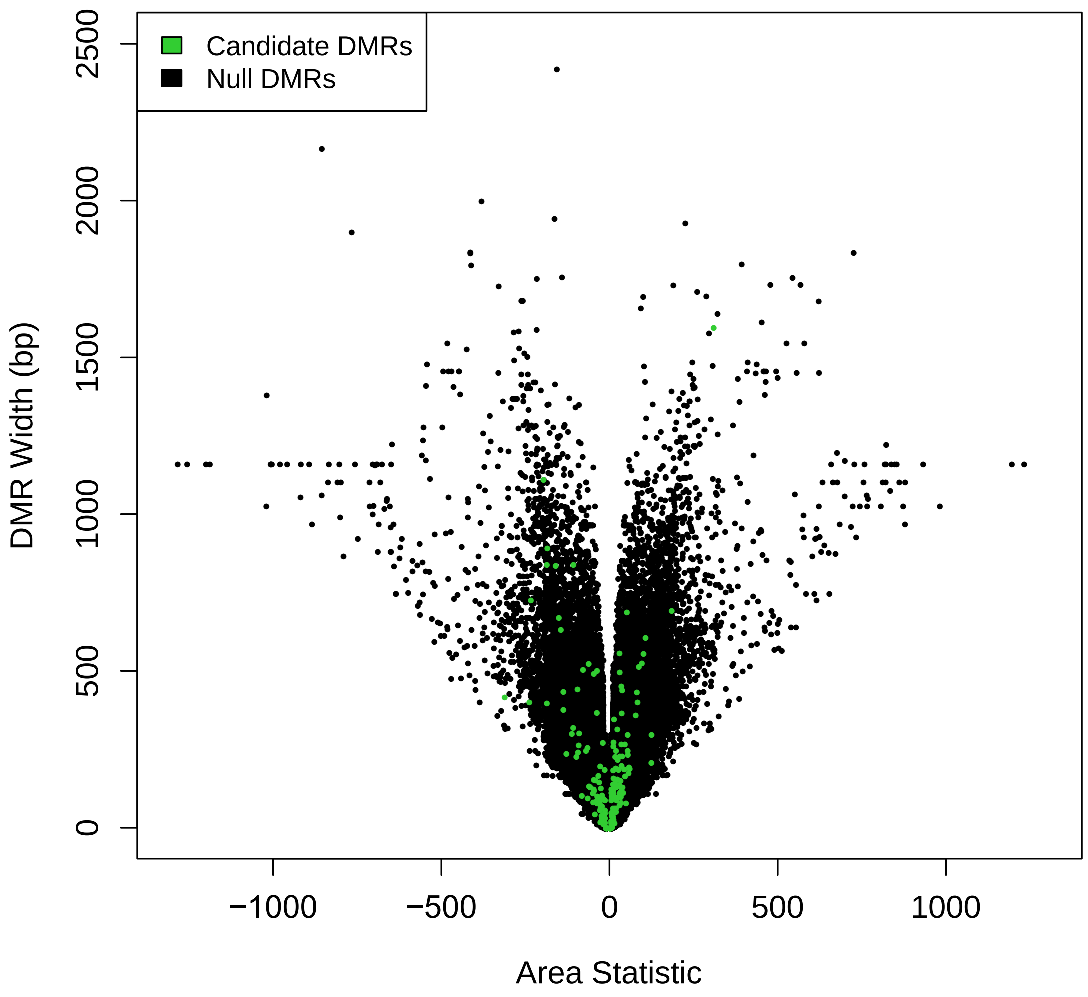

**Supplementary Figure S1:** Genome-wide results for identification of differentially methylated regions in placenta associated with fetal sex. Y-axis depicts width (bp) of null (black dots) or candidate (green dots) DMRs identified via BSmooth. X-axis depicts area statistics (sum of t-statistics in DMR) of null or candidate DMRs. Null DMRs are those generated across 1000 permutations.

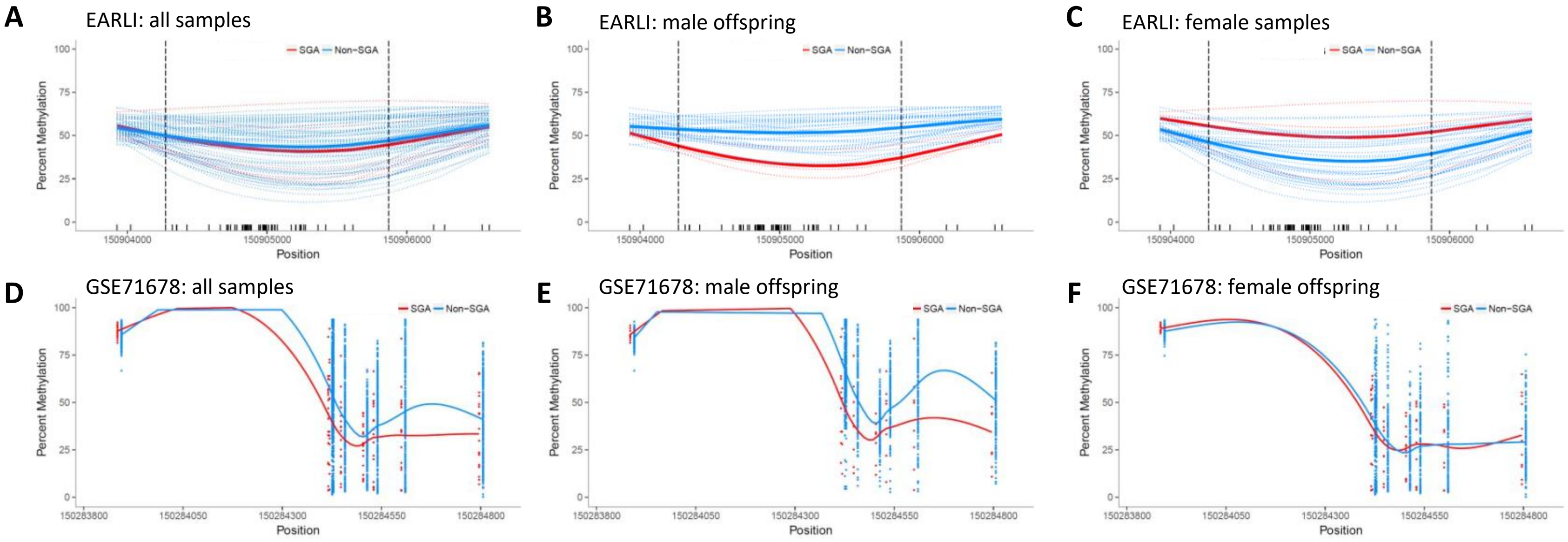

**Supplementary Figure 2: Inverse relationship between *ZNF300* promoter methylation and small for gestational age (SGA) between males and females.** Percent methylation (y-axis) vs genomic position (x-axis). Small for gestational age (SGA) is denoted in red and non-SGA in blue. EARLI study placenta whole genome bisulfite sequencing data (WGBS) from **A,** all samples (n=74); **B,** male offspring (n=34); **C,** female offspring (n=34). Illumina 450K DNA methylation data from replication set 3 (GSE71678) for **D**, all samples (n=338); **E**, male offspring (n=180); **F,** female offspring (n=158)

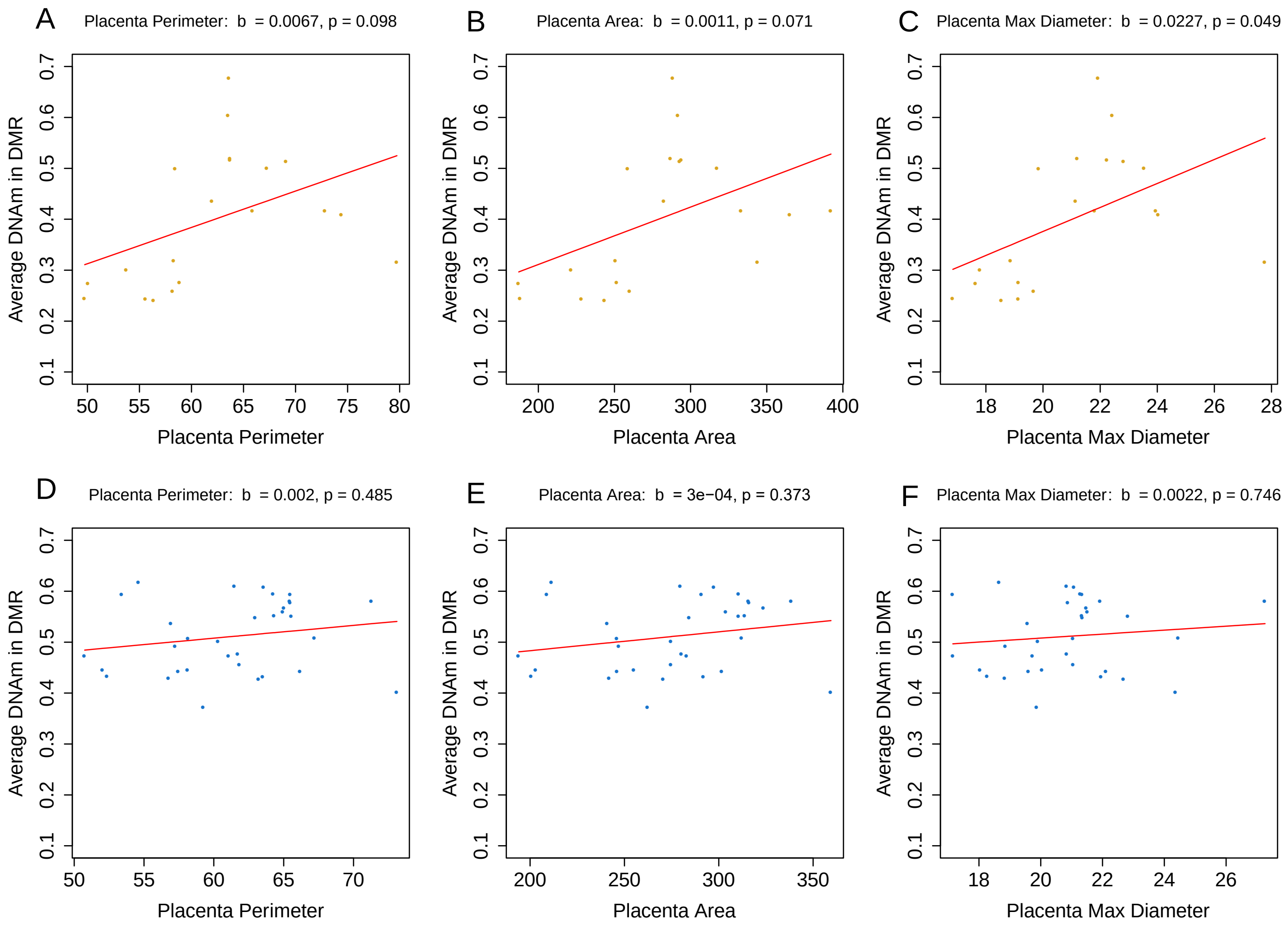

**Supplementary Figure 3: DNA methylation associations with placenta morphological features at *ZNF300* locus are observed in females but not males.** Mean methylation levels in DMR (y-axis) vs. placental perimeter (A,D), area (B,E), and maximum diameter (C,F), on the x-axis. P-values shown are for the regression of average methylation in the DMR onto each placental feature, adjusted for gestational age and FPR. A-C, among females (n=20; yellow points); D-F, among males (n=33; blue points).

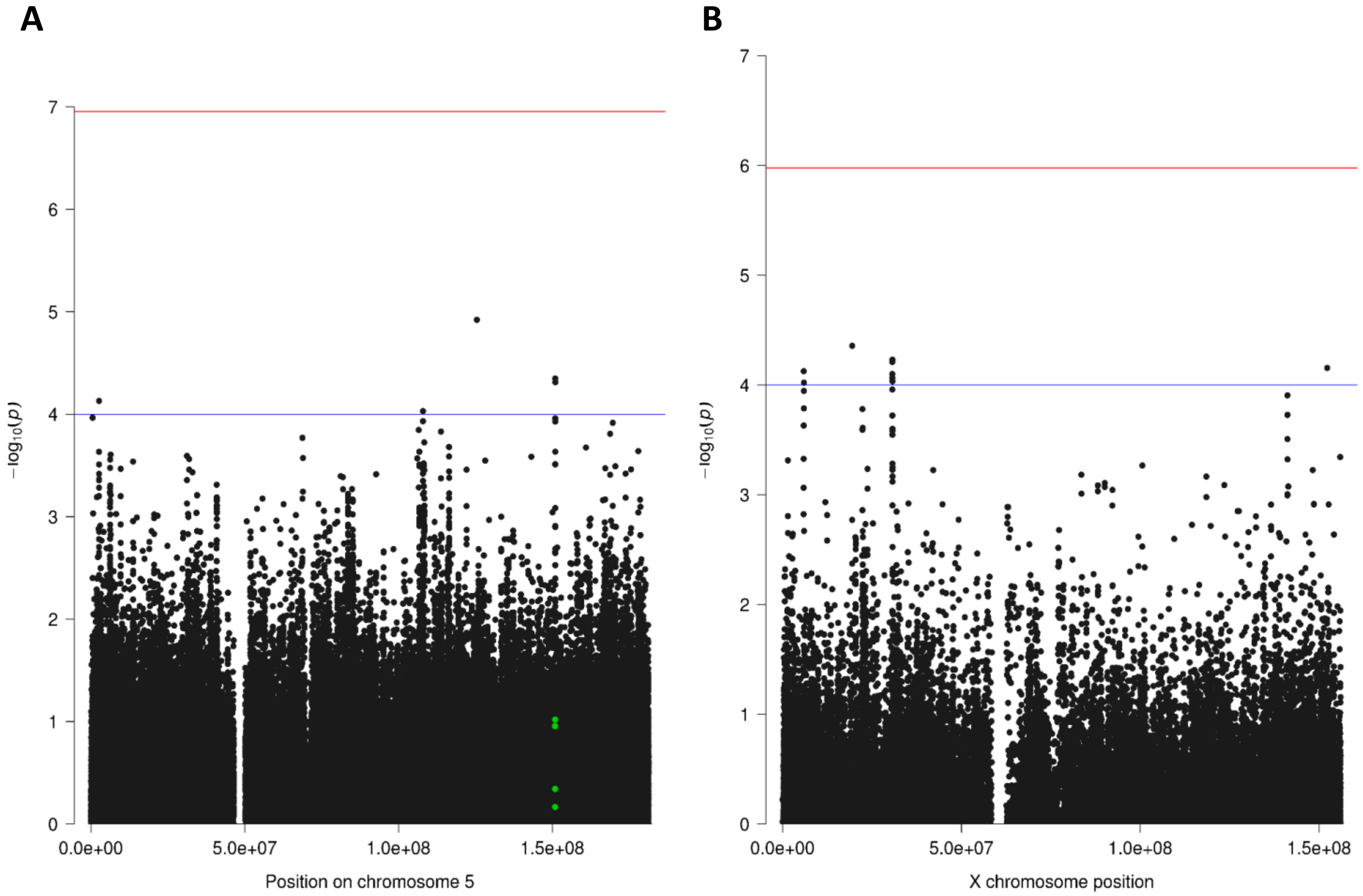

**Supplementary Figure 4: Manhattan plots showing SNP associations with DNA methylation at *ZNF300*. A,** chromosome 5 SNPs with green dots indicating SNPs that are located within the *ZNF300* differentially methylated region; **B,** SNPs on chromosome X that showed suggestive associations with methylation at *ZNF30*, in *trans*

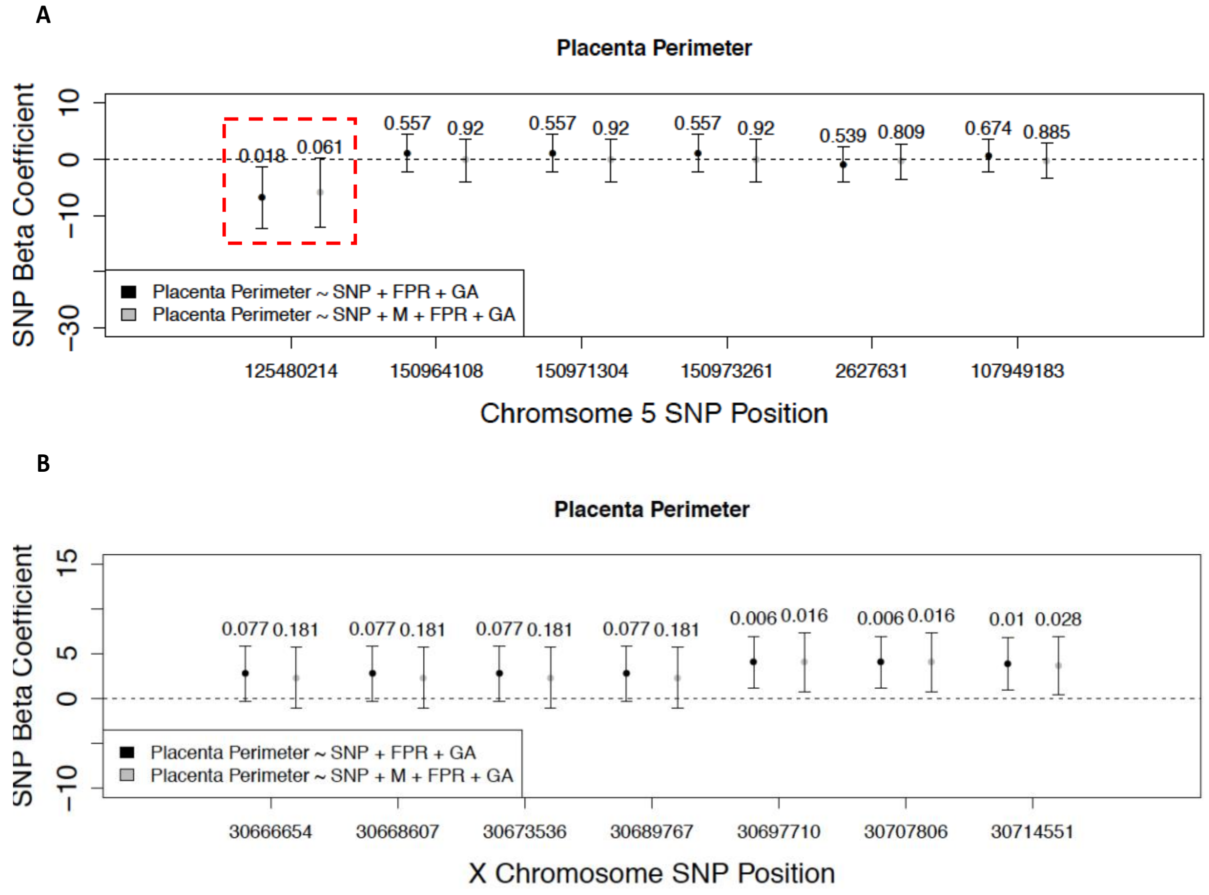

**Supplementary Figure 5: Causal inference testing reveals mediation of genetic variation on placenta perimeter by DNA methylation at the *ZNF300* locus.** Points indicate regression coefficients for SNP genotype terms in a model with the morphological phenotype as an outcome regressed onto the SNP, feto-placental weight ratio (FPR) and gestational age (GA) without (black) and with (gray) average methylation in the *ZNF300* DMR region. Corresponding p-values and error bars are also shown for each point. Red squares are drawn around SNPs showing evidence for mediation of SNP effects on placenta perimeter by *ZNF300* methylation.A**)** Results for chromosome 5 SNPs, and **B)** results from chromosome X SNPs.

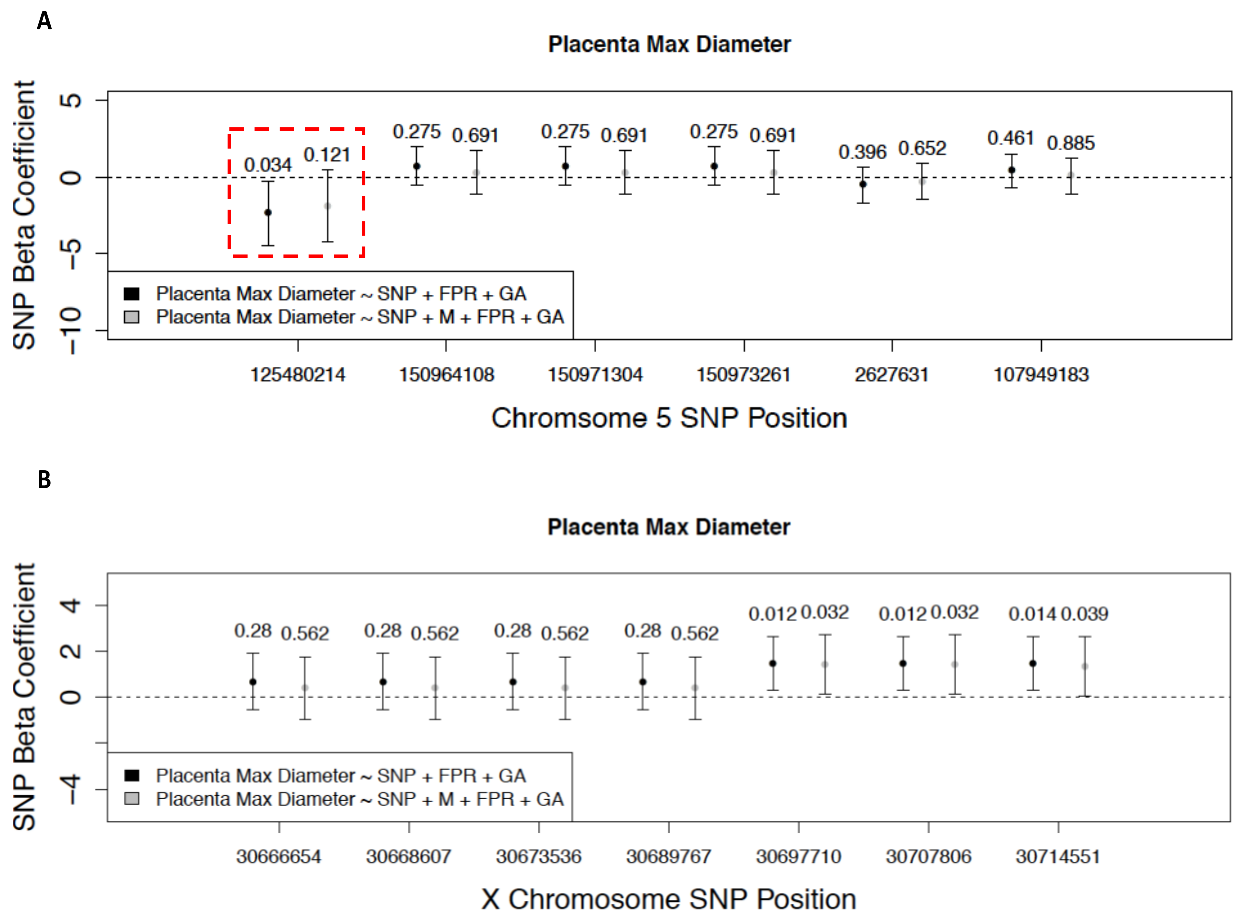

**Supplementary Figure 6: Causal inference testing reveals mediation of genetic variation on placenta max diameter by DNA methylation at the *ZNF300* locus.** Points indicate regression coefficients for SNP genotype terms in a model with the morphological phenotype as an outcome regressed onto the SNP, feto-placental weight ratio (FPR) and gestational age (GA) without (black) and with (gray) average methylation in the *ZNF300* DMR region. Corresponding p-values and error bars are also shown for each point. Red squares are drawn around SNPs showing evidence for mediation of SNP effects on placenta max diameter by *ZNF300* methylation. **A)** Results for chromosome 5 SNPs, and **B)** results from chromosome X SNPs.

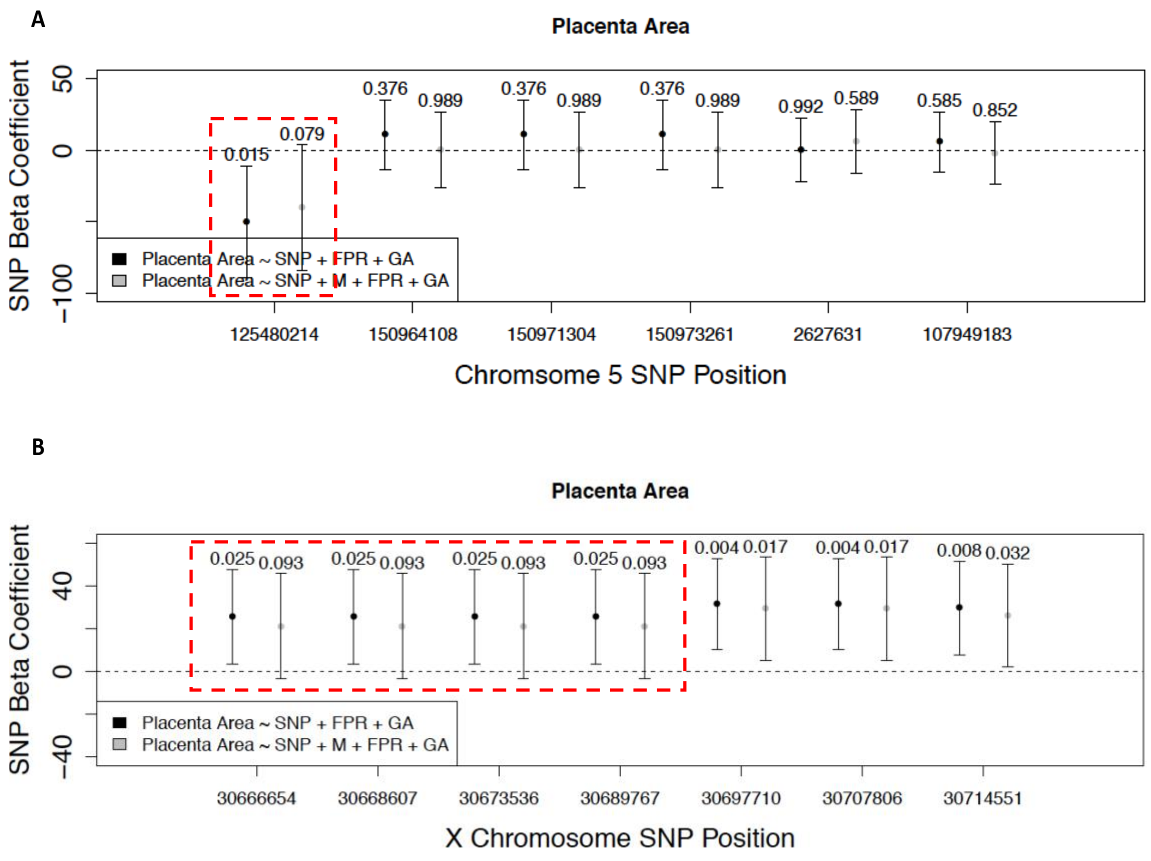

**Supplementary Figure 7: Causal inference testing reveals mediation of genetic variation on placenta area by DNA methylation at the *ZNF300* locus.** Points indicate regression coefficients for SNP genotype terms in a model with the morphological phenotype as an outcome regressed onto the SNP, feto-placental weight ratio (FPR) and gestational age (GA) without (black) and with (gray) average methylation in the *ZNF300* DMR region. Corresponding p-values and error bars are also shown for each point. Red squares are drawn around SNPs showing evidence for mediation of SNP effects on placenta area by *ZNF300* methylation. **A)** Results for chromosome 5 SNPs, and **B)** results from chromosome X SNPs.
